# Neural mechanisms underlying robust target selection in response to microstimulation of the oculomotor system

**DOI:** 10.1101/2024.10.29.620929

**Authors:** Mohsen Rakhshan, Robert J Schafer, Tirin Moore, Alireza Soltani

**Author notes:** **Corresponding author:** AS, Department of Psychological and Brain Sciences, Dartmouth College, Hanover NH 03755,.

## Abstract

Despite its prevalence in studying the causal roles of different brain circuits in cognitive processes, electrical microstimulation often results in inconsistent behavioral effects. These inconsistencies are assumed to be due to multiple mechanisms, including habituation, compensation by other brain circuits, and contralateral suppression. Considering the presence of reinforcement in most experimental paradigms, we hypothesized that interactions between reward feedback and microstimulation could contribute to inconsistencies in behavioral effects of microstimulation. To test this, we analyzed data from electrical microstimulation of the frontal eye field of male macaques during a value-based decision-making task and constructed network models to capture choice behavior. We found evidence for microstimulation-dependent adaptation in saccadic choice, such that in stimulated trials, monkeys’ choices were biased toward the target in the response field of the microstimulated site (*T_in_*). In contrast, monkeys showed a bias away from *T_in_* in non-stimulated trials following microstimulation. Critically, this bias slowly decreased as a function of the time since the last stimulation. Moreover, microstimulation-dependent adaptation was influenced by reward outcomes in preceding trials. Despite these local effects, we found no evidence for the global effects of microstimulation on learning and sensitivity to the reward schedule. By simulating choice behavior across various network models, we found a model in which microstimulation and reward-value signals interact competitively through reward-dependent plasticity can best account for our observations. Our findings indicate a reward-dependent compensatory mechanism that enhances robustness to perturbations within the oculomotor system and could explain the inconsistent outcomes observed in previous microstimulation studies.

## Introduction

Electrical microstimulation of the brain is one of the most widely used methods in systems neuroscience for uncovering the contributions of different brain circuits to cognition and behavior (Bradley et al., 2005; Clark et al., 2011; Hanks et al., 2006a; Murphey & Maunsell, 2007; Schafer & Moore, 2007; Seidemann et al., 1998). In addition, microstimulation has been extensively used for cognitive therapy (Grover et al., 2023) and to treat neurological diseases such as Parkinson’s and epilepsy (Wu et al., 2021) and to develop prosthetic devices for visual (Davis et al., 2012; Schmidt et al., 1996; Torab et al., 2011), somatosensory (Berg et al., 2013; K. Ding et al., 2024; Hughes et al., 2021; Tabot et al., 2013; Thomson et al., 2013; Urdaneta et al., 2021), and motor systems (Penfield & Boldrey, 1937; Sironi, 2011; Zimnik & Churchland, 2021). Although focused on different brain circuits and applications, one of the most direct tests for the effectiveness of microstimulation is its causal effects on choice behavior.

However, the influence of microstimulation on behavior is often inconsistent and associated evidence inconclusive, as divergent behaviors emerge across different experimental settings and animal models and even effects change over extended periods within the same study (Chamberlin & Saper, 1994; Elsner et al., 2016; Griffin et al., 2009; Hanks et al., 2006b; Murasugi et al., 1993; Plow et al., 2009; Salzman et al., 1992; Watanabe & Munoz, 2010, 2011). This inconsistency has been hypothesized to be due to many factors, including differences in characteristics and protocols of the microstimulation (Murasugi et al., 1993), technological and experimental design limitations (Doty, 1969; Jazayeri & Afraz, 2017), habituation, and compensation by the corresponding brain areas in the opposite hemisphere (Doty, 1969). Interestingly, these inconsistencies can also be viewed as robustness of behavior to microstimulation. Nevertheless, it is relatively easy to induce a behavioral shift locally (in time) when microstimulation is applied.

For example, previous studies that have used electrical microstimulation of different structures within the oculomotor system have found that microstimulation generates a bias in target selection toward the response field of the stimulated neurons (superior colliculus (SC) (Carello & Krauzlis, 2004), frontal eye field (FEF) (Gold & Shadlen, 2000; Murd et al., 2020; Schafer & Moore, 2007), lateral intraparietal cortex (LIP) (Hanks et al., 2006b), supplementary eye field (SEF) (Berdyyeva & Olson, 2014), and a combination of SC and FEF (Gardner & Lisberger, 2002). Critically, most of the tasks used in these studies involve reward feedback to motivate appropriate behavioral responses. Therefore, the overall behavior might remain robust to microstimulation due to interactions between reward feedback and microstimulation. Interestingly, activity in most areas of the oculomotor system is modulated by the reward value of the upcoming saccade including SC (Ikeda & Hikosaka, 2003), FEF (L. Ding & Hikosaka, 2006), LIP (Platt & Glimcher, 1999; Seo et al., 2009; Sugrue et al., 2004), and SEF (Chen & Stuphorn, 2015; Donahue et al., 2013). Therefore, the local behavioral bias caused by microstimulation of the above brain areas may interact with reward-value signals, and this could happen within the same areas.

Similarly, although a meta-analysis of studies on cognitive therapy using electrical stimulation indicates that such interventions can enhance cognitive functions such as attention and memory, the effects on motor learning and decision making remain inconclusive (Grover et al., 2023). Interestingly, both motor learning and decision-making tasks often involve reward feedback on a trial-by-trial basis. This suggests that inconsistency in the effectiveness of electrical stimulation for these tasks might be due to the interaction between stimulation and reward feedback.

To examine the interaction between reward feedback and electrical stimulation and its impact on learning and choice behavior, we analyzed data from microstimulation of the FEF during a dynamic value-based choice task in which monkeys chose between two alternative options that provided rewards with different probabilities. We compared the effects of microstimulation on choice behavior between trials in which microstimulation occurred, the subsequent trials with no microstimulation, and on average. Additionally, to capture our experimental findings and reveal possible neural mechanisms underlying the interaction between reward signals and microstimulation and its effect on target selection, we constructed multiple alternative network models of subcortical and cortical circuits within the oculomotor system to simulate choice behavior under different experimental conditions.

## Materials and Methods

### Experimental design and statistical analysis

Two male monkeys (Macaca mulatta) weighing 6 kg (monkey 1) and 11 kg (monkey 2) were used in the experiments. All surgical and behavioral procedures were approved by the Stanford University Administrative Panel on Laboratory Animal Care and the consultant veterinarian and were in accordance with the National Institutes of Health and Society for Neuroscience instructions. More details on the experimental design, including stimulus presentation, data acquisition, analysis of eye movements, and delivery of electrical stimulation are reported elsewhere (Schafer & Moore, 2007). Some non-overlapping findings based on this dataset have been published previously (Schafer & Moore, 2007; Soltani et al., 2021).

#### Visual stimuli

Saccade targets were drifting sinusoidal gratings within stationary, 5–8^°^ Gaussian apertures. Gratings had Michelson contrast between 2%-8% and a spatial frequency of 0.5 cycle/s^°^. Drift speed was 5°*/*s in a direction (up or down) perpendicular to the saccade required to acquire the target (left or right). These parameters were held constant during a session of the experiment. Targets were identical on each trial except for the drift direction (i.e., up versus down) selected randomly for each target.

#### The choice task with a dynamic reward schedule

Following fixation on a central fixation spot, there was a variable delay (200–600 *ms*) before the two targets appeared on the screen (**Fig. 1A**). Targets appeared simultaneously, equidistant from the fixation spot, and opposite to each other. Using a similar convention for the control and microstimulation conditions, we call these two targets *T_in_* and *T_out_*. In the control sessions (i.e., those without FEF microstimulation) *T_in_* and *T_out_* were in the left and right hemifields, respectively. In the microstimulation sessions, *T_in_* refers to the target within the FEF response field (within the left hemifield for both monkeys), while *T_out_* was the opposite. If the monkey directed a saccadic eye movement into an 8° invisible window around either target within 400 *ms* of the target appearance, it was delivered a juice reward according to the dynamic reward schedule described below.

**Figure 1.**
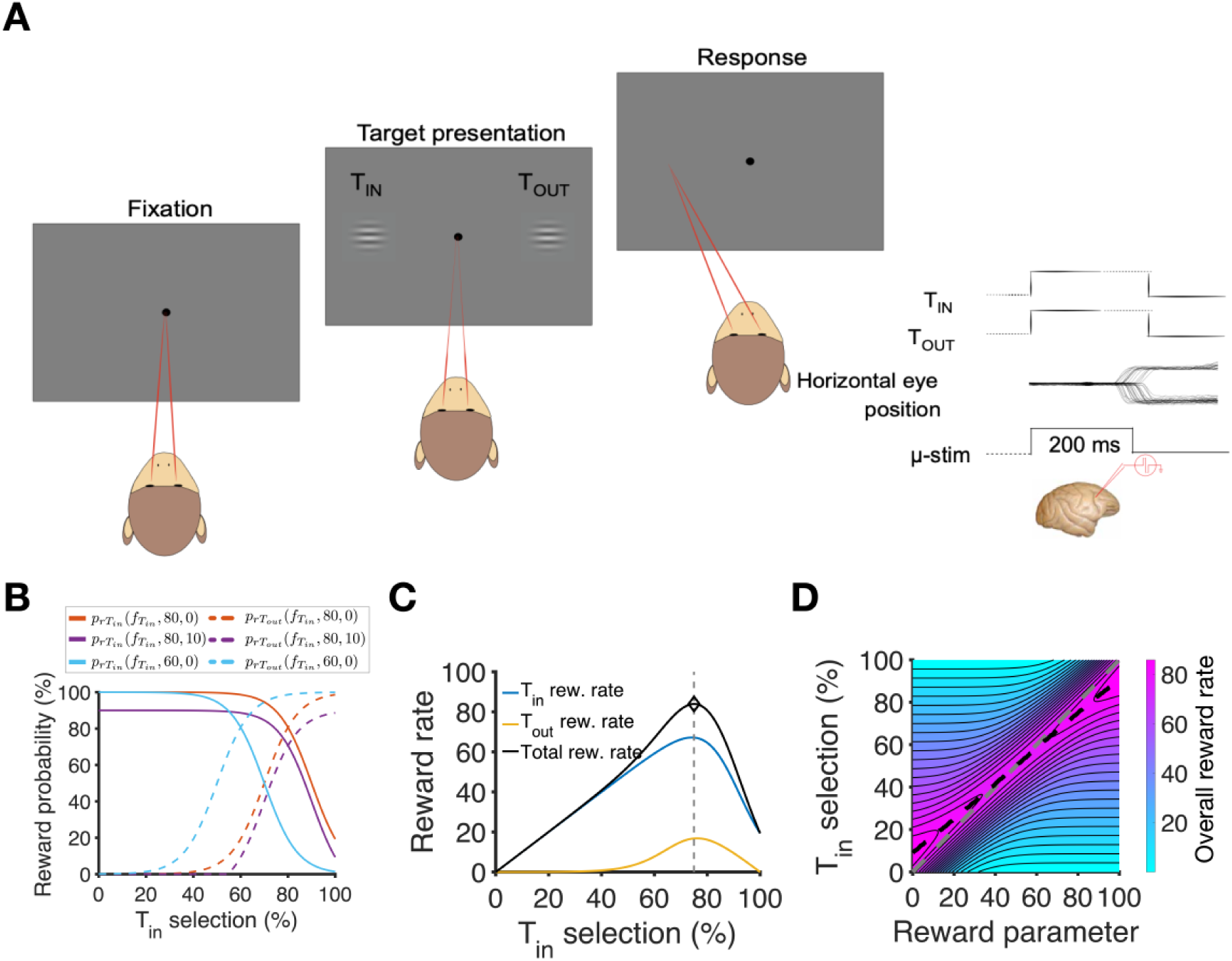
Dynamic value-based choice task and subthreshold microstimulation of the FEF. (**A**) Task design. A fixation point appeared on the screen on each trial, followed by the presentation of two drifting-grating targets. Monkeys indicated their choice with a saccade to one of the two targets; however, the targets were removed as soon as the eyes left the fixation point. A juice reward was delivered on a variable schedule following each saccade. *T_in_* and *T_out_* refer to the visual targets at the center of and directly opposite the RF of the FEF stimulation site, respectively. Event plots indicate the visual targets’ presentation sequence, saccade examples, and microstimulation signal and timing. The dashed lines denote variable time intervals. (**B**) Example reward schedules. Plot shows three examples of reward probability on the 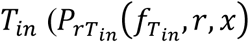, solid lines) and *T_out_* targets 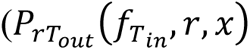, dashed lines) as a function of the local percentage of *T_in_* selection (*f*_*T_in_*_). The reward probability changes based on global reward parameter *r*, penalty parameter *x*, and the local *T_in_* selection percentage *f*_*T_in_*_ (**Equations 1–2**). (**C**) Reward harvest rate as a function of the local percentage of *T_in_* selection (*f*_*T_in_*_) for *r*=80 and *x*=0. (**D**) Reward harvest rate as a function of global reward parameter *r* and local *T_in_* selection percentage *f*_*T_in_*_. The gray dashed line corresponds to matching for which *f*_*T_in_*_ = *r*. The black dashed line shows the behavior that results in the optimal reward rate. The difference between the gray and black dashed lines indicates that a slight undermatching will result in optimal choice behavior in the task.

Overall, two monkeys completed 32 microstimulation sessions (19 and 13 sessions for monkeys 1 and 2, respectively) consisting of 7693 trials (4554 and 3139 trials for monkeys 1 and 2, respectively). They also completed 160 sessions of the dynamic choice task with no microstimulation (control condition; 74 and 86 sessions for monkeys 1 and 2, respectively). Each session consisted of an average of 140 and 370 trials for monkeys 1 and 2, respectively. The control and microstimulation conditions were performed in separate sessions.

#### Reward schedule

Following each correct saccadic choice, the monkey received a juice reward according to a dynamic schedule (Abe & Takeuchi, 1993). The probability of reward given a selection of *T_in_* or *T_out_* was equal to:

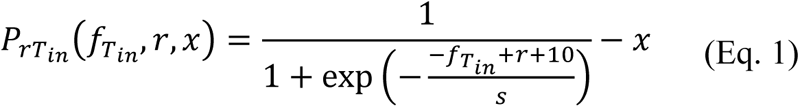

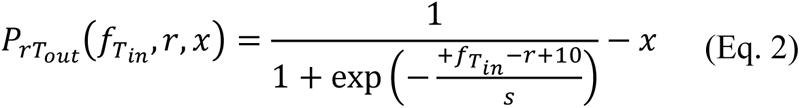

where *f_T_in__* is the local fraction of *T_in_* selection (in percentage) estimated using 20 trials preceding the current trial (**Fig. 1C**). Furthermore, *r*, the global reward parameter, is a task parameter that was held constant on a given session of the experiment. It dictates which option is more likely to yield a reward (*T_in_* for *r>50*, *T_out_* for *r<50*) when the two targets are selected with equal frequency (i.e., at *f_T_in__* = 50). *s* is another task parameter (set to 7 in all experiments) that determines the extent to which the deviation from the optimal strategy reduces reward probability. *x* is a penalty parameter used to adjust the global reward rate, which was kept constant throughout each session. Positive values of *x* decreased reward probability on saccades to both *T_in_* and *T_out_* targets to motivate monkeys further to identify and choose the more rewarding location at the time. It was assigned to one of the following values in a fraction of sessions as reported in the parentheses: 0 (77%), 0.15 (6%), 0.30 (6%), or 0.40 (11%) in the control condition, and 0 (53%), 0.05 (6%), 0.10 (3%), or 0.15 (38%) in the microstimulation condition. Due to the penalty parameter *x* and the configuration of reward schedule, 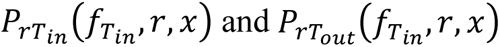 are not necessarily complementary. Lastly, any possible negative reward probability was substituted with 0 to prevent the reward probability from becoming negative.

Based on the configuration of the reward schedule, the reward probabilities on choosing *T_in_* and *T_out_* are equal at *f_T_in__* = *r*. This condition corresponding to matching behavior is slightly suboptimal in this task, as shown in **Figure 1D, E** (see (Soltani et al., 2021) for more details). As the value of *s* approaches zero, matching and optimal behavior become closer to one another. As shown in our previous work (Soltani et al., 2021), we found that the performance of both monkeys was suboptimal.

#### Electrical microstimulation

Electrical stimulation of an FEF site was delivered through tungsten electrodes while current amplitude was measured via the voltage drop across a 1 *kOhm* resistor in series with the return lead of the current source. First, the FEF was localized on the basis of its surrounding physiological and anatomical landmarks and the ability to evoke fixed-vector, saccadic eye movements with (suprathreshold) stimulation using currents below 50 *µA* at a frequency of 200 *Hz* (0.3 *ms* pulse duration, 100 *ms* trains). Threshold current was determined using a separate calibration task. During the stimulated trials of the choice task, subthreshold microstimulation was delivered at threshold current (±2 *µA*) at a lower frequency of 60 *Hz*. Thresholds for evoking saccades were measured again after the experimental sessions, and sessions with significantly different current thresholds were excluded from analyses.

The location of the endpoints of evoked saccades due to suprathreshold microstimulation was then expressed as the center of the FEF site’s response field (RF). To target sessions in which microstimulation was efficacious but did not simply drive the monkey’s eyes toward one target, we analyzed sessions in which microstimulation changed the probability of *T_in_* selections by a number between 0% and 20%, as described in Schafer and Moore (Schafer & Moore, 2007).

During sessions of saccadic choice trials that included microstimulation (microstimulation condition), target *T_in_* was presented at the center of the RF and *T_out_* was opposite. Microstimulation was delivered on one-half of the randomly assigned trials. Subthreshold microstimulation was achieved by reducing the frequency (and number) of current pulses to 60 *Hz*, from 200 *Hz*, while holding the current constant. This was done as an alternative to reducing the microstimulation current (Moore & Fallah, 2001) to minimize the change in the stimulated volume of the cortex between threshold and subthreshold conditions while greatly altering its efficacy (Gardner & Lisberger, 2002). Microstimulation trains began at the same time as the appearance of the targets and lasted for 200 *ms*.

#### Statistical analysis

Statistical tests used for comparing different conditions are reported in the text, along with the p-value and effect size. Unless otherwise specified, the two-sided Wilcoxon sign test or Wilcoxon signed-rank test is used to compare paired data, whereas the two-sided Wilcoxon rank-sum test is used to compare independent data.

### Reinforcement learning models used for fitting choice data

We used various reinforcement learning (RL) models to fit choice behavior in control and microstimulation conditions. The details for all the RL models are previously reported (Soltani et al., 2021). We found that choice behavior in control and microstimulation sessions can best be explained using the same RL model. In this RL model, the locations of the *T_in_* and *T_out_* targets in trial *t* were assigned with subjective reward values (related to reward probability) of 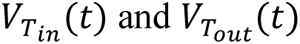, respectively. These subjective reward values were updated at the end of each trial based on the reward outcome of the trial and the learning rule described below. Furthermore, we used a sigmoid function of the difference in subjective reward values of the two targets to compute the probability of selecting *T_in_*:

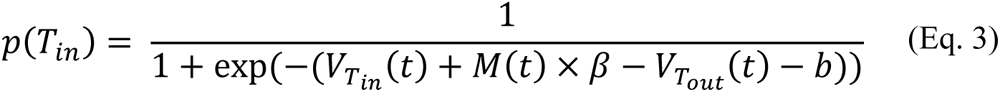

where 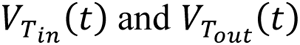 are the subjective reward values for the *T_in_* and *T_out_* targets, respectively, *M*(*t*) is a variable indicating whether microstimulation is applied in trial *t* (i.e., *M*(*t*) = 1 for a microstimulated trial (MS+) and *M*(*t*) = 0 for a non-microstimulated trial (MS-)), and β is a coefficient biasing the choice towards *T_in_* target in MS+ trials. Finally, *b* is the general bias in choice behavior toward the *T_in_* target to capture any side bias.

At the end of each trial, the subjective reward values of both chosen and unchosen targets were updated based on the reward outcome of that trial. More specifically, while subjective reward values of both chosen and unchosen targets were discounted in the following trial, the value of chosen target was further updated depending on the reward outcome. For example, if *T_in_* was selected in trial *t* and the trial was rewarded (or not rewarded), the subjective reward values were updated as follows:

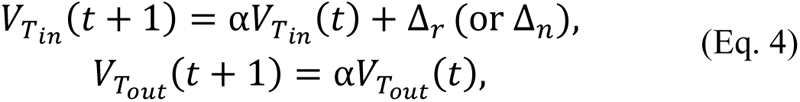

where 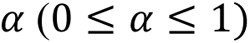 is the discount factor determining how much the estimated subjective reward value from one trial carries over to the following trial and Δ_*r*_(Δ_*n*_) determines the change in subjective reward value if the trial was rewarded (not rewarded). Therefore, discount factors closer to 1 correspond to a longer-lasting effect of reward (i.e., integration of reward on longer timescales) or slower learning.

### Network models

Multiple brain areas involved in saccadic choice also receive dopaminergic input (including the FEF), and the interaction between these areas determines adaptive target selection. To model this interaction in the context of FEF microstimulation, we considered a model that includes a second target-selection circuit in addition to the FEF circuit. We only included two target-selection circuits to avoid the complexity associated with the interaction between three or more circuits. Therefore, the network model consisted of a total of three circuits: the valuation circuit corresponding to a circuit estimating reward probability and two target-selection circuits corresponding to two areas in the oculomotor system.

The valuation circuit consisted of two pools of value-encoding neurons that were activated upon the presentation of visual targets and projected to the corresponding pools of neurons in the target-selection circuits. The reward value (or reward probability) associated with each target was encoded in two sets of plastic synapses onto the value-encoding pools. At the end of each trial, these neurons received signals indicating whether *T_in_* or *T_out_* target was selected and whether a reward was delivered. Their afferent plastic synapses were modified using a stochastic, reward-dependent, Hebbian learning rule (Soltani & Wang, 2006) (see next section). This modification enabled plastic synapses to estimate the reward value of each target, which in turn was reflected in the activity of the value-encoding pools.

The two target-selection circuits simulated two cortical areas involved in saccadic target selection, one of which was the FEF that was the site of microstimulation in our experiment. For simplicity of the modeling, we considered similar properties for the two target-selection circuits. Each target-selection circuit consisted of two pools of excitatory neurons selective for the two targets and one non-selective pool of inhibitory neurons. The selective pools received excitatory inputs from the corresponding value-encoding pools in the valuation circuit and inhibitory input from a shared inhibitory pool of neurons within each circuit. This architecture enabled each circuit to select between the two targets based on the reward value of the two targets (Soltani & Wang, 2006). Since the sensory inputs to value-encoding neurons were similar, the only factor differentiating the inputs to decision neurons was the strength of the plastic synapses from the sensory neurons onto the value-encoding neurons. As has been shown before, the selection behavior of each decision circuit can be fit as a sigmoid function of the difference in the inputs (Soltani et al., 2006; Soltani & Wang, 2006, 2010)

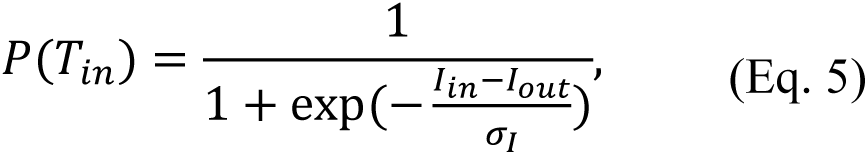

where *P*(*T_in_*) is the probability of selecting *T_in_*; *I_in_* and *I_out_* are the input currents from the value-encoding neurons, and *σ*_*I*_ (in mA) describes the sensitivity of the target-selection circuit to biased inputs (*I_in_* − *I_out_*). Therefore, instead of running the firing-rate network on each trial, we used the inputs from the valuation circuit into each target-selection circuit to obtain choice probability using **Equation 5**. We then used the combination of the outcome choice probabilities based on the two target-selection circuits to determine the selection of the model in each trial (see below).

To simulate the microstimulation experiment, we assumed that the pool of decision neurons selective for *T_in_* in the FEF target-selection circuit received an extra input on stimulated trials, which mimicked microstimulation currents. In addition, we assumed that the input from the *T_in_* value-encoding pool to the corresponding decision neurons was plastic (see below).

### Learning reward probabilities in the valuation circuit

The input currents from the value-encoding neurons depend on factors including the synaptic strength, total number of plastic synapses in each neuron, presynaptic firing rate, peak conductance of the potentiated and depressed states, and the time constant of input synaptic current (Soltani et al., 2006; Soltani & Wang, 2006). However, except for the synaptic strength, all other factors are similar for the two value-encoding pools and could be factored into a single variable k (in *mA*). Therefore, the selection behavior can be expressed as a function of the difference in the synaptic strengths of the neurons encoding the reward probability for *T_in_* and *T_out_* targets (i.e., *c_in_* and *c_out_*), respectively:

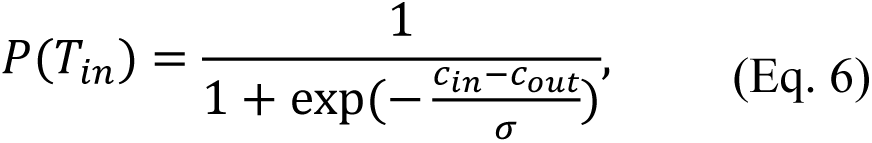

where 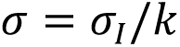 is normalized sensitivity.

At the end of a given trial, according to the trial’s choice and reward outcome, the synaptic strengths for *T_in_* and *T_out_* targets (*c_in_* and *c_out_*, respectively) were modulated according to the following rules. If *T_in_* target was chosen and rewarded, *c_in_* was increased, but *c_out_* remained intact:

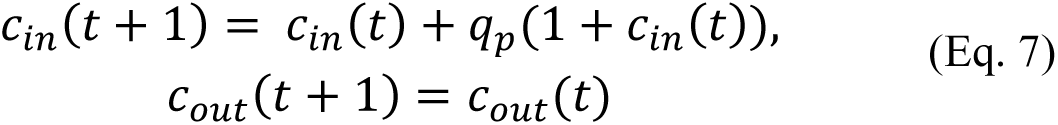

where *q_p_* is the potentiation constant and resembles the learning rate of rewarded trials in a reinforcement learning model (set to 0.15 in our simulations). In contrast, if *T_in_* target was chosen, but not rewarded, *c_in_* was decreased, but *c_out_* remained the same:

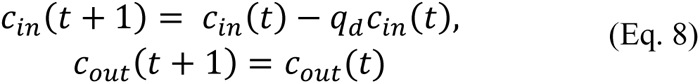

where *q_d_* is the depression constant and resembles the learning rate of unrewarded trials in a reinforcement learning model (set to 0.15 in our simulations). Similarly, if *T_out_* target was chosen and rewarded, *c_out_* was increased, but *c_in_* remained intact:

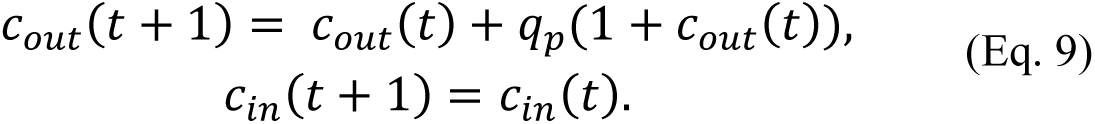

In contrast, if *T_out_* target was chosen but not rewarded, *c_out_* was decreased, but *c_in_* remained the same:

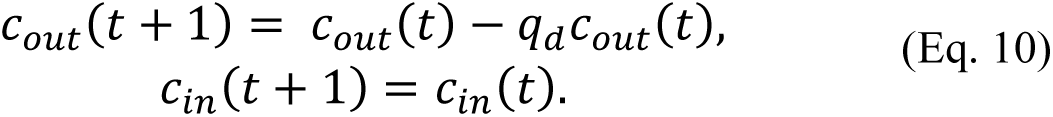

### Alternative mechanisms for the effects of microstimulation

Here, we constructed three general models for the effects of microstimulation on target selection. This exploration was done for two main reasons. First, the effects of microstimulation on neural activity are very complicated processes that depend on the sensitivities of different neural elements (i.e., dendrites, axons, initial segments) as well as the properties of single neurons and the connectivity of neurons within the stimulated network (Histed et al., 2009; Tehovnik et al., 2006). Secondly, our goal was not to capture the effect of microstimulation on neural activity but to find general mechanisms that result in the observed patterns of adaptation of microstimulation effects. Therefore, to simulate our experimental observations and identify the most plausible mechanism, we started with the least complex model (i.e., the model with no adaptation to microstimulation). Based on the simulation results, we progressively increased the model’s complexity (details below).

#### The model with no adaptation to microstimulation

In this most basic model, the effect of the microstimulation on decision making was simulated as a constant extracellular input current, *I*_μ_, that affected the *T_in_* pool in the first target-selection circuit (in the FEF) with efficacy *k*_μ_. This assumption was based on general observations that stimulation of a site in the oculomotor system bias the saccades toward the object placed in the receptive field of the stimulated site (Carello & Krauzlis, 2004; Gold & Shadlen, 2000; Murd et al., 2020; Schafer & Moore, 2007). Therefore, we assumed that neurons in the FEF *T_in_* target-selection pool received an additional current equal to *k*_μ_ *I*_μ_, resulting in the probability of *T_in_* selection in the FEF target-selection circuit to be equal to:

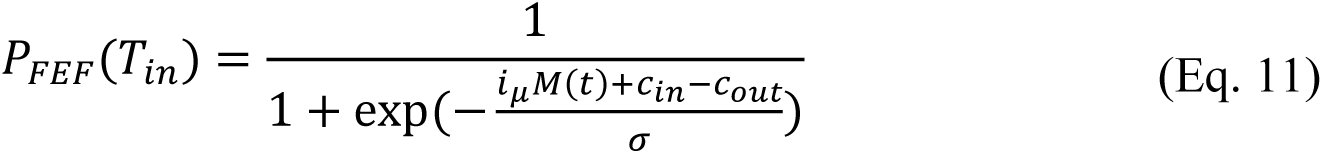

where *I*_μ_ = *k*_μ_ *I*_μ_/*k* is a dimensionless quantity that determines the ratio of the strength of the microstimulation current to the reward input currents, and *M*(*t*) encodes the presence or absence of the microstimulation on trial *t*; it was equal to 1 if there was microstimulation and 0 otherwise.

The second target-selection circuit received the same input currents from the value-encoding neurons as the FEF target-selection circuit. However, to simulate cross-area compensatory mechanisms in the oculomotor system, we assumed that microstimulation in the FEF accompanied by an increase and a decrease in input to the neurons selective for *T_in_* and *T_out_* targets in the second target-selection circuit. This compensation mechanism was based on the results of two studies by Schlag and colleagues (Schlag et al., 1998; Schlag-Rey et al., 1992), where they identified three groups of neurons based on their reactions to microstimulation in the contralateral FEF: 35% of the neurons were unaffected, 37% showed inhibition, and 20% exhibited excitation. Crucially, they found that the neurons that became excited had response selectivity similar to that of the stimulated FEF neurons, whereas the neurons that became inhibited had response selectivity opposite to that of the stimulated FEF neurons.

Based on this compensatory mechanism, we set the probability of *T_in_* selection in the second target-selection (STS) circuit equal to:

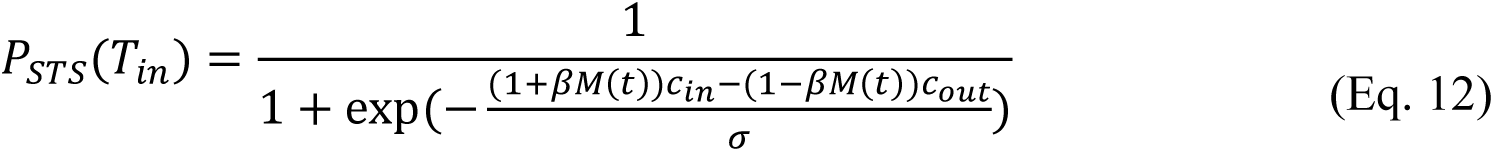

where β is a dimensionless factor that increase and decrease the value-dependent input to *T_in_* and *T_out_* pools in STS, respectively (set to 0.2 × *i*_μ_ in our simulations; see **Equation 11**).

Finally, the outcome choice probabilities of the two target-selection circuits are combined to determine the final choice probability for selection between the two targets, *P*(*T_in_*), on each trial using the following equation:

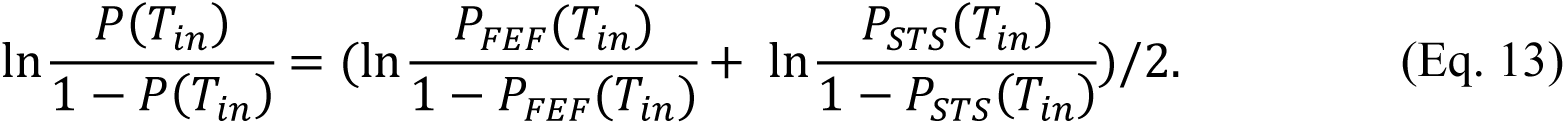

#### Model with desensitization

In this model, microstimulation affected the *T_in_* pool in the FEF target-selection circuit similarly to the previous model (i.e., the base model with no adaptation to microstimulation), but in addition, we assumed that due to stimulation, the neurons selective for *T_in_* within the FEF target-selection circuit became less sensitive to their inputs. This assumption was based on the evidence showing that neurons in the central nervous system exhibit “fatigue” (Ye et al., 2012) or an increased refractory period as a response to stimulation (Feng et al., 2014). However, rather than modeling the ion channels in detail to simulate fatigue or changes in the refractory period, we simulated these effects by simply including depression or desensitization in the inputs to *T_in_* neurons (Dayan & Abbott, 2001).

Specifically, after each microstimulation, the depression variable *d(t)* was updated as follows:

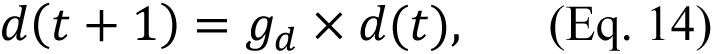

where 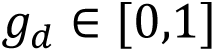 (set to 0.9 in our simulations) determines the percent change in sensitivity due to synaptic depression. In the absence of microstimulation, the depression variables decayed back to 1 using the following equation:

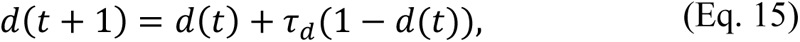

where τ_*d*_ is the time constant which determines the dynamics of adaptation (set to 0.85 in our simulations). Therefore, the probability of *T_in_* selection in the FEF target-selection circuit was equal to

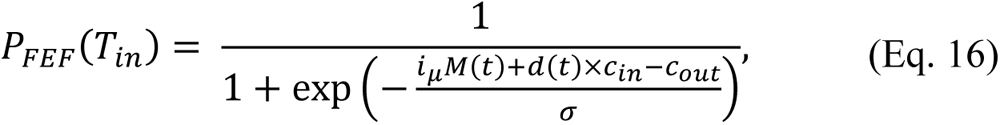

where, similar to the model with no adaptation to microstimulation, *i*_μ_ is a dimensionless quantity that determines the ratio of the strength of the microstimulation current to the reward input currents (set to 0.5 in our simulations). Finally, the second target-selection circuit was modeled similarly to the model with no adaptation (**Equation 12**), and the outcome choice probabilities of the two target-selection circuits were combined using **Equation 13**.

#### The model with reward-dependent adaptation

This model was built upon the previous model (i.e., the model with desensitization) with one difference: in this model, the efficacy of the microstimulation current, *k*_μ_(*t*), varied from trial to trial in the FEF target-selection circuit. This efficacy depended on the state of the individual neuron and the network architecture (i.e., the architecture of the network of neurons close to the electrode tip and their connections to the rest of the neurons in the FEF *T_in_* target-selection pool). We did not directly model these factors directly; instead, we assumed that during the microstimulation experiment, individual neurons in the FEF *T_in_* target-selection pool adapted through processes akin to homeostasis, maintaining consistent activity levels. Given the incomplete understanding of such homeostatic plasticity mechanisms, we modeled this effect by adjusting the efficacy of microstimulation in each trial based on reward outcomes.

More specifically, we assumed this efficacy consisted of two components: a fixed component that depended on network dynamics and a variable component capturing the combination of network and single-cell adjustments to microstimulation:

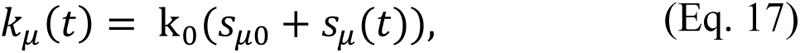

where *s*_μ0_ is the fixed component (set to 1.5 in our simulations) and *s*_μ_(*t*) is the variable component, and *k*_0_ is a constant determining the overall efficacy. As a result, the FEF *T_in_* target-selection pool received an additional current equal to *k*_μ_(*t*)*I*_μ_.

To maintain the same level of activity, the efficacy of *I_in_* input to the *T_in_* target-selection pool, *k_in_*, was modulated by the following factors through two different mechanisms:

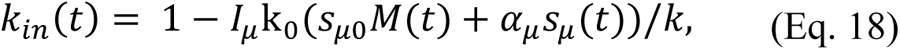

where *I*_μ_ determines the ratio between the effects of two types of adjustments on *I_in_* efficacy and α_μ_ determines the relative strength of adjustments in input current due to changes in microstimulation current efficacy (set to 1.5 in our simulations). Using **Equation 5**, the choice probability in the FEF target-selection circuit can be written as

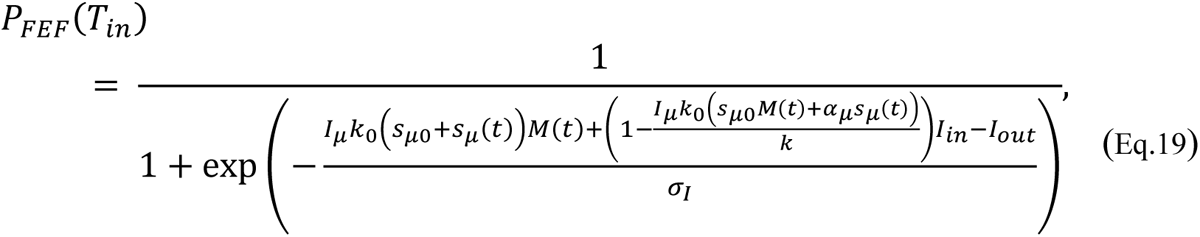

and by normalizing by the common factors for the reward input currents, *k*, we obtain

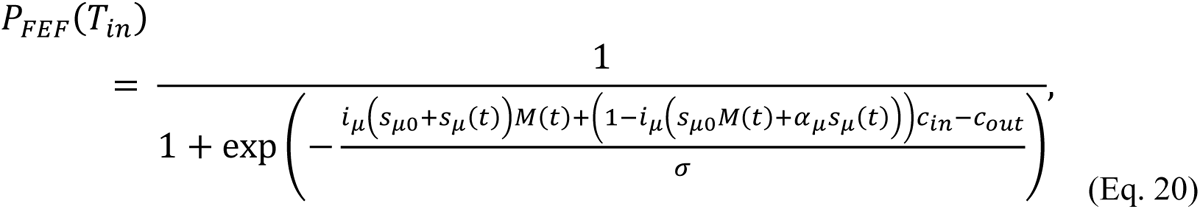

where *i*_μ_ = *k*_0_*I*_μ_/*k* is a dimensionless quantity which determines the ratio of the strength of the microstimulation current to the reward input currents (set to 0.7 in our simulations).

Moreover, the adaptive part of the microstimulation current efficacy in the FEF circuit was updated as

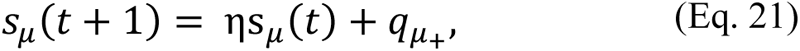

if microstimulation occurred and the choice was rewarded, where η is a discount factor (set to 0.5 in our simulations) and *q*_μ+_ quantifies the change in efficacy of microstimulation after a rewarded trial (set to 0.05 if *T_in_* target was chosen and 0.95 if *T_out_* target was chosen). On the other hand, if the choice was not rewarded or there was no microstimulation in the trial, this quantity was discounted

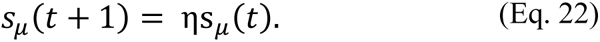

The second target-selection circuit was modeled similarly to those in the desensitization model and base model with no adaptation.

#### Models without oculomotor system interaction

To determine whether our experimental data could be replicated by models with a single target-selection circuit, we simulated three such models even though these models do not align with existing knowledge of the oculomotor system. These include a base model with no adaptation to microstimulation, a model with desensitization, and a model with reward-dependent adaptation. These network models consisted of two circuits: the valuation circuit, which corresponded to a circuit estimating reward probability, and one target-selection circuit, which corresponded to the FEF. The valuation circuit and the target-selection circuit were similar to the circuits for the network models considering cross-area compensation (see above). We made three models based on the model’s adaptation to microstimulation similar to the models with cross-area compensation (see Alternative mechanisms for the effects of microstimulation above for details). These three models were the base model with no adaptation to microstimulation, the model with desensitization, and the model with reward-dependent adaptation.

### Estimation of confidence

To estimate monkeys’ confidence on a given trial, we employed two methods. In the first method, we used the reaction time of each trial as a proxy for confidence, a method commonly used in the literature (Martino et al., 2013; Stolyarova et al., 2019). The trials with faster reaction times were associated with higher confidence, whereas those with slower reaction times indicated lower confidence. Therefore, we used the median reaction time from each session to classify trials into high- and low-confidence categories depending on whether a trial had a reaction time faster or slower than the median reaction time, respectively.

In the second method, we defined high- and low-value confidence trials based on the difference between the values of chosen and unchosen targets, as estimated by the best-fitting RL model. Again, we used the median difference between the values of targets of each trial (chosen-unchosen) to divide trials into high- and low-confidence trials based on whether this difference was smaller or larger than the median difference. The rationale behind this categorization was that a larger difference between the values of the two targets increases the likelihood that the higher-value target would be chosen, corresponding to easier decision making and, thus, higher confidence. We performed this analysis using estimated values from the RL model rather than actual reward values because these estimates reflect subjective reward expectations, making them more relevant to confidence assessments. Because both methods yielded qualitatively similar results and reaction time is more commonly used to estimate confidence, we have opted to present only the results based on this method.

## Results

### Local effects of microstimulation on saccadic choice

To examine whether microstimulation of the oculomotor system and reward feedback interact, we analyzed data from a dynamic value-based decision-making task in which monkeys were trained to select between one of the two targets that appeared on the screen simultaneously. The monkeys freely selected between the two targets using saccadic eye movements under control and microstimulation conditions (**Fig. 1A**). In the control sessions, following the selection of a target, the monkey received a juice reward according to a dynamic schedule that was a function of the local fraction of choosing that target in the past 20 trials. In each session, one target was globally more rewarding, but this reward decreased every time the monkey chose that target (**Equations 1–2**; **Fig. 1C–E**). Therefore, to successfully perform this task, the monkeys had to consider choice and reward history to estimate the reward probability associated with the two targets.

The microstimulation sessions were identical to the control ones except that in half of the randomly selected trials, subthreshold microstimulation was delivered at the site of the FEF, beginning when the targets were presented and lasting for 200 *ms*. Accordingly, one of the two targets (*T_in_*) was placed within the response field (RF) of the FEF. The other target (*T_out_*) was placed in the opposite hemifield (**Fig. 1B**; see **Materials and Methods** for details). The choice behavior of monkeys, its optimality, and the reinforcement learning analyses in the control condition––which occurred in separate sessions from the microstimulation condition––have been previously described (Soltani et al., 2021). Here, we compared the behavior of the monkeys in the control and microstimulation conditions to examine how microstimulation of the FEF affected behavior.

To test the local effects of microstimulation on target selection, we divided trials in the microstimulation condition into those with and without microstimulation (MS+ and MS-trials, respectively). We found that microstimulation biased target selection on a trial-by-trial basis. Specifically, we found that monkeys chose *T_in_* target more often in the MS+ trials than in the MS-trials (Wilcoxon sign test; two monkeys together: median(difference) = 0.066; *p* = 2.5 ∗ 10^−7^, *d* = 1.26, monkey 1: median(difference) = 0.081; *p* = 7.3 ∗ 10^−4^, *d* = 1.36, monkey 2: median(difference) = 0.040; *p* = 2.4 ∗ 10^−4^, *d* = 1.46). Furthermore, even though the microstimulation was subthreshold and did not evoke saccades by itself, it increased the probability of choosing *T_in_* in microstimulated trials (MS+ trials) compared to the control condition (Wilcoxon sign test; two monkeys together: median(difference) = 0.038; *p* = 2.5 ∗ 10^−7^, *d* = 0.89; monkey 1: median(difference) = 0.035; *p* = 7.3 ∗ 10^−4^, *d* = 0.96; monkey 2: median(difference) = 0.033; *p* = 2.4 ∗ 10^−4^, *d* = 1.03; **Fig. 2**). In addition, the probability of *T_in_* selection in MS-trials was smaller than the probability of *T_in_* selection in the control condition (Wilcoxon ranksum test, two monkeys together: median(difference) = -0.085; *p* = 0.0015, *d* = 0.23, monkey 1: median(difference) = - 0.1; *p* = 7.9 × 10^−4^, *d* = 0.35, monkey 2: median(difference) = -0.073; *p* = 0.12, *d* = 0.15).

**Figure 2.**
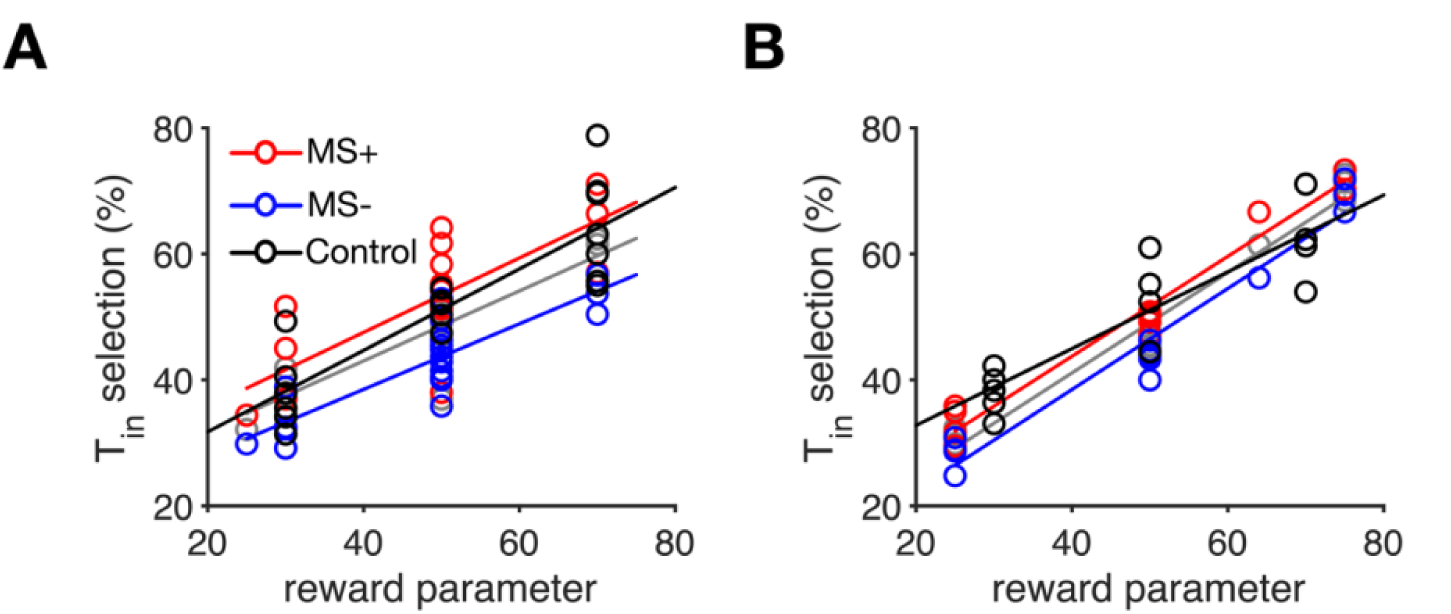
Overall target selection and changes in saccadic choice in response to the FEF microstimulation. The percentage of *T_in_* selections in each session is plotted as a function of the global reward parameter in that session for the control condition (black symbol), for all trials of the microstimulation condition (gray symbols), and separately for stimulated (MS+: red symbols) and non-stimulated trials (MS-: blue symbols) of the microstimulation condition. Results for monkey 1 and monkey 2 are plotted in panels (A) and (B), respectively. The solid lines show linear fits.

Despite increased and decreased *T_in_* selection on MS+ and MS-trials, the overall *T_in_* selection in the microstimulation condition was not significantly different than the average *T_in_* selection in the control condition (Wilcoxon ranksum test, two monkeys together: *p* = 0.06, *d* = 0.13, monkey 1: *p* = 0.05, *d* = 0.21, monkey 2: *p* = 0.24, *d* = 0.12; **Fig. 2**). Lack of evidence for an overall bias toward *T_in_* in the microstimulation condition, despite clear effects of microstimulation on a trial-by-trial basis, suggests a potential compensatory mechanism within the oculomotor system and its related circuits. We explore this possibility further below.

**Figure 3.**
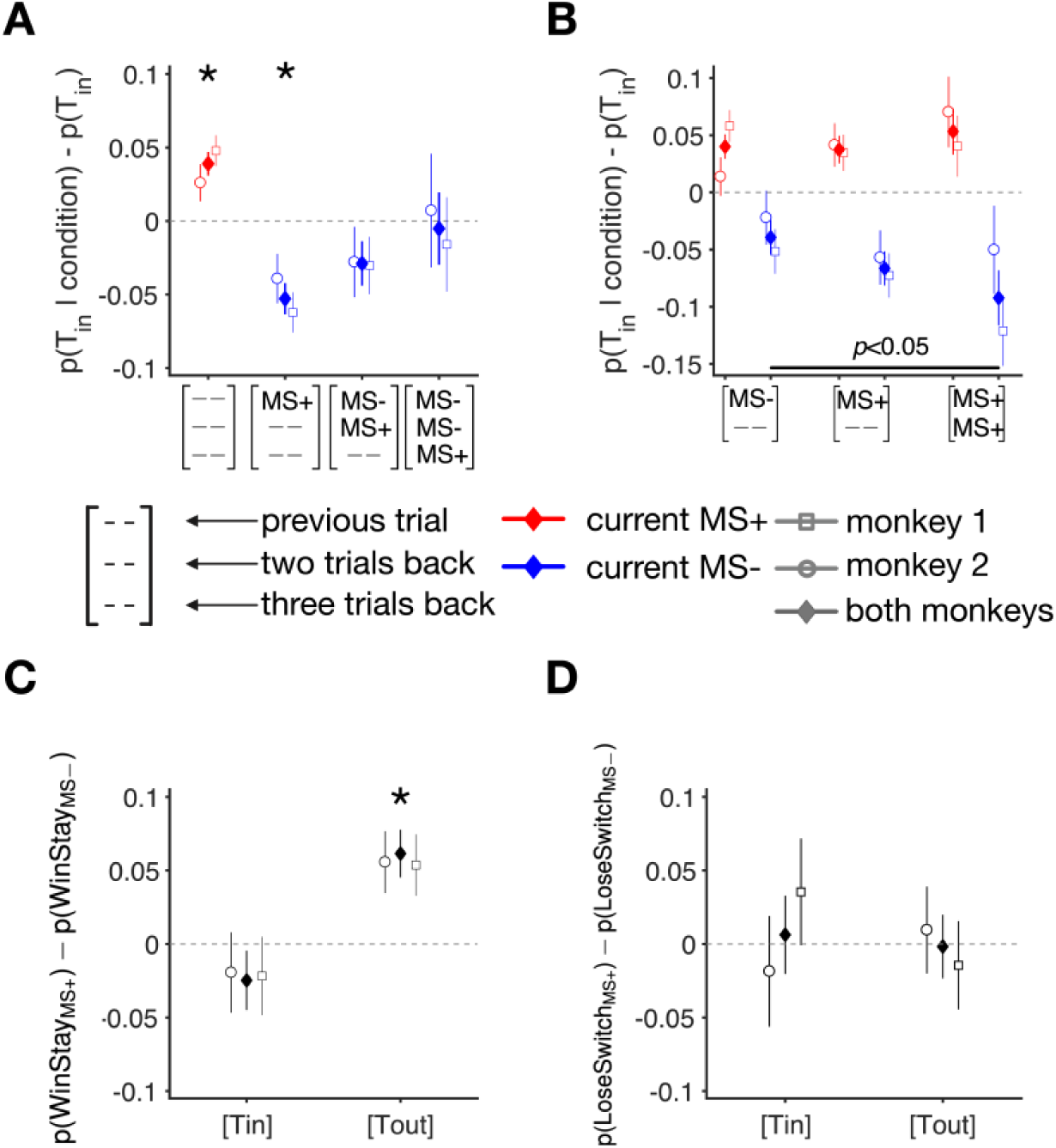
Evidence for compensatory mechanisms for microstimulation effects and interactions between microstimulation and reward feedback. (**A**) Change in *T_in_* selection on MS+ trials (red) and on MS-trials (blue) that occurred one, two, or three trials following an MS+ trial. Average values for monkey 1 (square) and monkey 2 (circle), and both monkeys (diamond) are shown separately, and error bars show the s.e.m. Asterisks indicate values significantly different from zero (Wilcoxon sign test, *p<*0.001). The ‘--’ sign indicates that the corresponding trial could have been either MS+ or MS-. (**B**) Effects of consecutive microstimulation on target selection. Change in *T_in_* selection is plotted for different numbers of consecutive MS+ trials (red symbols) and on the following MS-trials (blue symbols). Symbols follow the same conventions as in panel A. (**C**) Differential response to reward in the presence and absence of microstimulation. The difference between the probability of stay after reward (*Win-Stay*) on MS+ trials and MS-trials is computed separately for when the monkey selected *T_in_* and *T_out_* on the preceding trial. Symbols follow the same conventions as in panel A. (**D**) Similar to panel C, but for the difference between the probability of switch after no reward (*Lose-Switch*) on MS+ and MS-trials.

To further examine the local effects of microstimulation on target selection, we evaluated the effect of microstimulation in non-stimulated trials following a microstimulated trial. To that end, we calculated the change in the probability of *T_in_* selection on MS-trials one, two, or three trials following an MS+ trial. We found that whereas microstimulation increased *T_in_* selection in the current trial (Wilcoxon sign test; two monkeys together: *p* = 2.5 ∗ 10^−7^, *d* = 0.89, monkey 1: *p* = 7.3 ∗ 10^−4^, *d* = 0.96, monkey 2: *p* = 2.4 ∗ 10^−4^, *d* = 1.03). However, this bias was reversed on the subsequent MS-trials (Wilcoxon sign test; two monkeys together: *p* = 1.1 ∗ 10^−4^, *d* = 0.71, monkey 1: *p* = 0.02, *d* = 0.73, monkey 2: *p* = 0.003, *d* = 0.79), and the trend dissipated within approximately three trials (**Fig. 3A**). This result points to microstimulation-dependent adaptation in saccadic choice.

To examine whether this microstimulation-dependent adaptation varied as a function of the number of consecutive microstimulations, we computed the change in *T_in_* selection for different numbers of consecutive MS+ trials and on their subsequent MS-trials (**Fig. 3B**). Although we did not find any evidence for an increase in *T_in_* selection as the number of consecutive MS+ trials increased (one-way ANOVA, *p* = 0.79), the bias away from *T_in_* selection on subsequent MS-trials was significantly increased (one-way ANOVA, *p* = 0.038). This result shows that the observed adaptation to microstimulation became stronger after consecutive microstimulation events.

We hypothesized that this microstimulation-dependent adaptation and the lack of overall bias towards *T_in_* in microstimulation sessions could be due to a possible interaction between microstimulation and reward feedback. This interaction could happen as bias caused by microstimulation interfered with how monkeys responded to the reward outcome in the preceding trials. We compared the differential response to reward feedback between the control and microstimulation conditions to test this hypothesis by computing the difference between the probability of *Win-Stay* and *Lose-Switch* strategies. *Win-Stay*, which involves choosing the same target as in the previous trial if it was rewarded, measures sensitivity to reward. *Lose-Switch*, which entails switching from the target selected in the previous trial if not rewarded, measures the sensitivity to the absence of reward. Therefore, the difference between the probability of *Win-Stay* and *Lose-Switch* strategies measures differential sensitivity to reward and absence of reward in which positive and negative values indicate greater sensitivity to the rewarded and unrewarded outcomes, respectively (Farashahi et al., 2017).

We found that the difference between the probability of *Win-Stay* and *Lose-Switch* was more negative for both monkeys in the microstimulation compared to the control condition (Wilcoxon ranksum test; both monkeys: Δ= −0.13, *p* = 1.1 ∗ 10^−4^, *d* = 0.28, monkey 1: Δ= −0.09, *p* = 0.08, *d* = 0.18, monkey 2: Δ= −0.12, *p* = 5.1 ∗ 10^−4^, *d* = 0.35). This result suggests a potential interaction between microstimulation and reward feedback, as monkeys showed higher sensitivity to unrewarded relative to rewarded outcomes during the microstimulation condition. Together, these findings point to mechanisms within the oculomotor systems that allow the effects of microstimulation on FEF to be compensated using reward feedback.

### Global effects of microstimulation on saccadic choice

To further test the hypothesis that the interaction of reward feedback and microstimulation could offset the local effects of microstimulation on saccadic choice, resulting in no overall effect, we assessed the global effects of microstimulation on overall task performance. First, we calculated the overall harvested reward as a function of the global reward parameter (*r*). Despite individual variations, we found no significant differences between the control and microstimulation conditions (**Fig. 4A, D**; **Fig. 5**). We also tested whether within the microstimulation sessions, microstimulation of the FEF changed the overall response of the monkeys to the reward schedule. To that end, we examined the slope of *T_in_* selection as a function of global reward parameter *r* during the microstimulation sessions (**Fig. 4B, E**). We did not find evidence that monkeys followed the reward schedule differently in MS+ and MS-trials (slope and confidence interval (*CI*): monkey 1: *m*_MS+_ = 0.59, *CI*(0.95)_MS+_ = [0.34,0.83], *m*_MS-_ = 0.52, *CI*(0.95)_MS-_ = [0.36,0.67]; monkey 2: *m*_MS+_ = 0.79, *CI*(0.95)_MS+_ = [0.70,0.88], *m*_MS-_ = 0.80, *CI*(0.95)_MS-_ = [0.69,0.92]).

We then compared overall target selection between the control and microstimulation conditions to examine the relationship between choice and reward fractions (i.e., matching behavior). Overall, we found that both monkeys followed the reward schedule similarly in the two conditions as reflected in the slope of choice vs. reward fractions (slope (*m*) of choice vs. reward fractions and its confidence interval: monkey1: *m*_MS_ = 0.55, *CI*(0.95)_MS_ = [0.40,0.71], *m*_control_ = 0.54, *CI*(0.95)_control_ = [0.46,0.62]; monkey2: *m*_MS_ = 0.80, *CI*(0.95)_MS_ = [0.71,0.88], *m*_control_ = 0.74, *CI*(0.95)_control_ = [0.71,0.76]; **Fig. 4C, F**).

**Figure 4.**
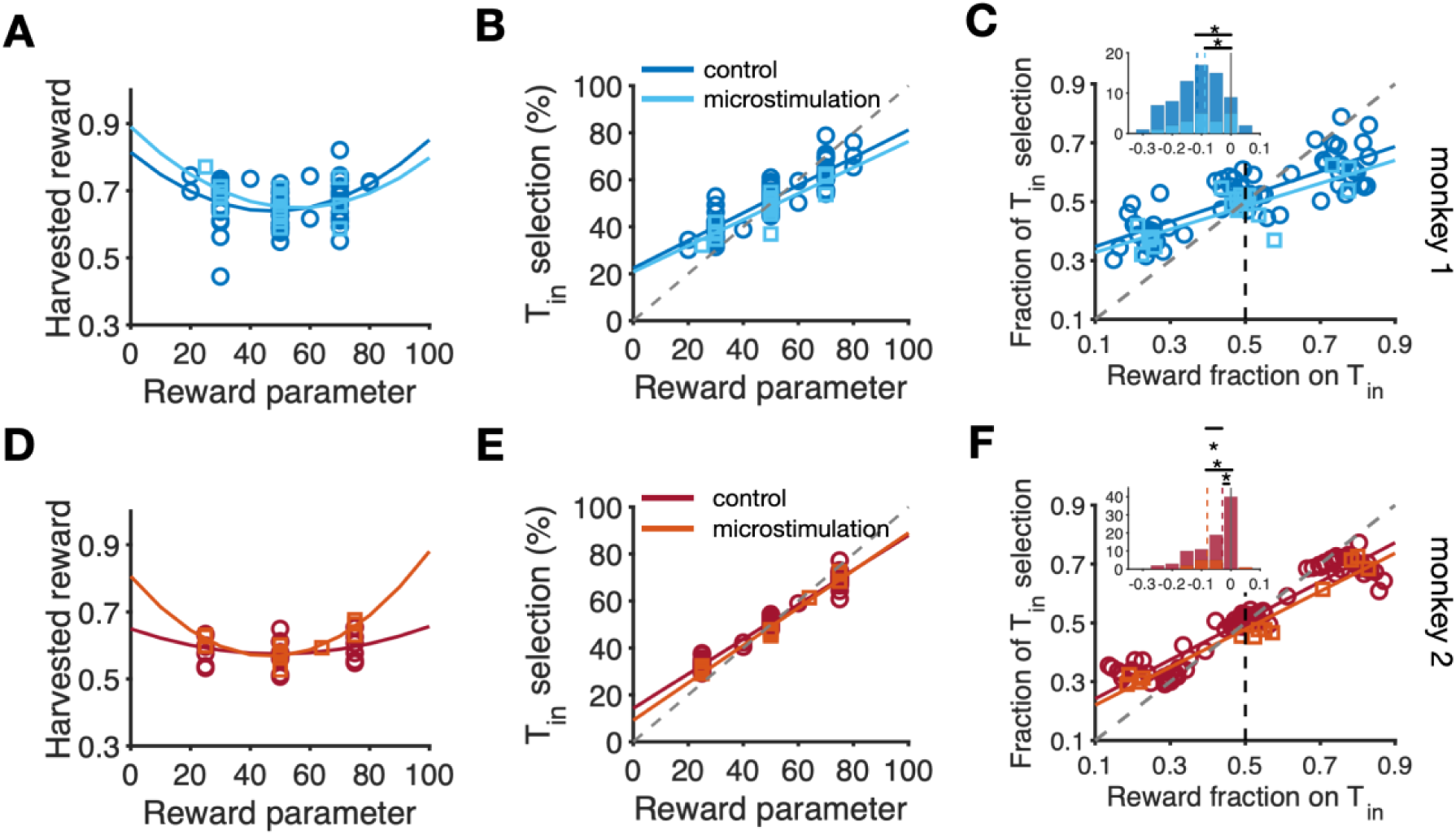
Overall target selection and changes in saccadic choice in response to the FEF microstimulation, separately for the two monkeys. (**A**) Harvested reward per trial as a function of the global reward parameter for zero penalty sessions for monkey 1. Dark (light) blue circles correspond to the control (microstimulation) sessions. The solid colored lines show fit using a quadratic function. (**B**) Percentage of *T_in_* selections as a function of the global reward parameter for the control (dark blue circle) and microstimulation sessions (light blue squares) of monkey 1. The gray dashed line is the diagonal line. The solid-colored lines show linear fits. (**C**) The proportion of *T_in_* selections as a function of the fraction of harvested reward on the *T_in_* target for monkey 1. The colored lines are linear fits. The gray dashed line shows the diagonal line corresponding to matching behavior. The inset shows the histogram of the difference between choice and reward fractions with negative and positive values corresponding to under- and over-matching. The dark and light blue histograms correspond to the control and microstimulation conditions. The colored dashed lines in the inset indicate the medians of the distributions, and asterisks show a significant difference from 0 (matching) using Wilcoxon signed-rank test (*p*<.05). (**D–F**) Similar to panels A–C, but for monkey 2. The results in panel D are shown for sessions with the global reward parameter equal to 15 (majority of the microstimulation sessions for monkey 2).

By comparing choice and reward fractions across sessions, we found that monkeys exhibited strong undermatching in both control and microstimulation conditions (**Fig. 4C, F**). That is, the relative selection of the more rewarding target was smaller than the relative reinforcement obtained on that target (monkey 1 control: median(choice fraction – reward fraction) = −0.115; Wilcoxon signed-rank test, *p* = 1.67 × 10^−12^, *d* = −1.35; monkey 1 microstimulation: median(choice fraction – reward fraction) = −0.0885; Wilcoxon signed-rank test, *p* = 3.98 × 10^−4^, *d* = −1.10; **Fig. 4C** inset; monkey 2 control: median(choice fraction – reward fraction) = −0.03, *p* = 1.76 × 10^−12^, *d* = −0.83; monkey 2 microstimulation: median(choice fraction – reward fraction) = −0.08, *p* = 4.88 × 10^−4^, *d* = −1.89; **Fig. 4F** inset). Crucially, the extent of undermatching did not differ significantly between the control and microstimulation conditions for monkey 1 (Wilcoxon ranksum test; *p* = 0.19, *d* = −0.14). However, monkey 2 exhibited slightly stronger undermatching in the microstimulation compared to the control condition (Wilcoxon ranksum test; *p* = 0.04, *d* = 0.21). These results point to either no change or slight reduced sensitivity to rewards due to stimulation.

**Figure 5.**
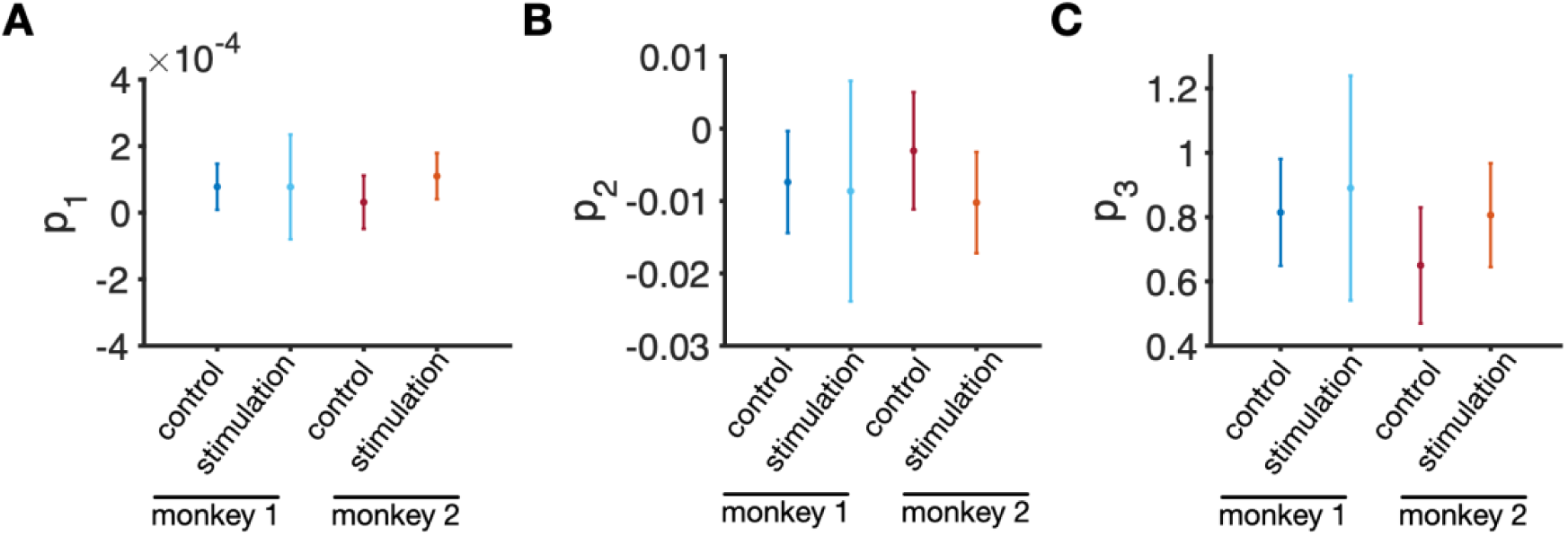
Comparison of the overall harvest rate between control and microstimulation conditions. Plotted are estimated values and confidence intervals for the overall harvested reward as a function of the global reward parameter using a quadratic function (harvested reward = *p*_1_*r*^2^ + *p*_2_*r* + *p*_3_). The error bars are the 98.75% confidence interval for each parameter. The confidence interval has been adjusted to 98.75% instead of the usual 95% to account for multiple comparisons (four comparisons across different conditions and monkeys).

### Effects of microstimulation on learning

To examine whether microstimulation influenced learning, we fit the choice behavior in control and microstimulation conditions using different reinforcement learning (RL) models. We found that a similar RL model best explained the choice behavior of both monkeys in the control and microstimulation sessions (see **Materials and Methods** for details). Interestingly, we did not find a significant difference between the discount factors of the RL model in the control and microstimulation conditions for either monkey (monkey 1: difference (stim-control) in discount factor ≈ 0; Wilcoxon ranksum test; *p* = 0.74, *d* = −0.03; monkey 2: difference (stim-control) in discount factor = 0.06; Wilcoxon ranksum test; *p* = 0.15, *d* = 0.14). Therefore, the comparison of discount factors, which indicates a difference in the learning rate, did not provide any evidence for an overall change in learning due to microstimulation (see **Materials and Methods** for details).

Together, the above analyses of monkeys’ choice behavior demonstrate that although microstimulation induced an overall bias in target selection on MS+ trials, there was no evidence that it impacted the monkeys’ ability to learn and perform the task effectively. These findings suggest that despite the local effects of FEF microstimulation on target selection, the oculomotor system and related circuits were able to compensate for these changes using reward feedback, thereby maintaining robust overall saccadic choice behavior in the face of perturbation.

### Neural mechanisms underlying adjustments of saccadic choice to microstimulation

To account for our observations, we constructed several computational network models to explore alternative plausible mechanisms. The basic model consists of three circuits: two target-selection circuits and a valuation circuit. The reward values of the two targets are encoded in two separate sets of plastic synapses in the valuation circuit. These synapses are updated at the end of each trial depending on the choice of the network and reward feedback. The two target-selection circuits represent two brain areas involved in saccadic choice: the FEF that was the site of microstimulation in our experiment and another area in the oculomotor system (e.g., LIP, SEF, etc.). Each target-selection circuit consists of two excitatory neuronal populations that are selective to the two targets and one non-selective inhibitory neuronal population. The excitatory pools of neurons in each target-selection circuit receive excitatory inputs from the corresponding pools of neurons in the value-encoding circuit and use these inputs to make decisions (see **Materials and Methods** for more details).

This configuration with two target-selection circuits allowed us to simulate the interaction between the FEF and another oculomotor area. We simulated this interaction based previous findings that microstimulation of FEF can cause both excitation and inhibition in the contralateral FEF (Schlag et al., 1998; Schlag-Rey et al., 1992).

Specifically, the contralateral neurons that became excited displayed response selectivity similar to that of the stimulated FEF neurons. In contrast, the neurons that were inhibited showed response selectivity that was opposite to that of the stimulated FEF neurons (see **Materials and Methods** for details).

To reveal the underlying mechanism of the observed effects of microstimulation on choice behavior, we implemented different mechanisms by which microstimulation could impact the behavior. To that end, we progressively increased the complexity of our models, starting from a base model with no adaptation to microstimulation, then to a model with desensitization to microstimulation, and finally, a model with reward-dependent adaptation to microstimulation (see **Materials and Methods** for more details).

A distinct effect of microstimulation at an oculomotor site is a biasing of saccades toward the object placed in the receptive field of the stimulated site (Carello & Krauzlis, 2004; Gold & Shadlen, 2000; Murd et al., 2020; Schafer & Moore, 2007). Therefore, we tested whether a model in which microstimulation increased input to the target in the receptive field of the stimulated site (*T_in_*) could replicate our observations. In this model, referred to as the model with no adaptation, microstimulation added a constant input current added to the *T_in_* pool of neurons in the FEF target-selection circuit. Moreover, to capture the aforementioned cross-area compensation in the oculomotor system, we assumed that an increase in input to *T_in_* pool in the FEF was accompanied by a decrease and increase in the value-dependent input to the neurons selective for *T_in_* and *T_out_* in the second target-selection circuit, respectively (see **Figure 6A** and **Materials and Methods** for details). This modulation of input to the second target-selection circuit helped to offset the increased input to the FEF caused by microstimulation.

We found that this model with no adaptation showed a decrease in *T_in_* selection on MS-trials following an MS+ trial. However, this effect was constant and did not decay to zero in the absence of microstimulation (**Fig. 6B**). Similarly, we found no significant effects of microstimulation repetition on target selection (**Fig. 6C**). Overall, these results demonstrate that within the proposed architecture, cross-area compensation through modulation of value-dependent input to the second target-selection circuit alone is insufficient to explain the observed adjustments in choice behavior, leading us to explore models with adaptation to microstimulation.

The first model with adaptation to microstimulation was based on the evidence showing that neurons in the central nervous system can exhibit “fatigue” (Ye et al., 2012) or an increased refractory period as a response to stimulation (Feng et al., 2014). We simulated these effects by including depression or desensitization in the inputs to T_in_ neurons. More specifically, in this model, referred to as the model with desensitization, the microstimulation caused a decrease in sensitivity of the FEF *T_in_* pool to its inputs (see **Figure 6A** and **Materials and Methods** for more details). This model had a similar cross-area compensation mechanism to the base model with no adaptation to microstimulation.

We found that the model with desensitization could account for the adaptation of the bias away from the *T_in_* target in MS-trials following a MS+ trial (**Fig. 6B**). However, the model also predicted a decrease in the effect of microstimulation across consecutive trials that, which was not observed in the experimental data (**Fig. 6C**). Thus, this result shows that the decrease in the sensitivity of the *T_in_* pool to its inputs from value-encoding circuit could not fully reproduce the experimental results using our dynamic model.

**Figure 6.**
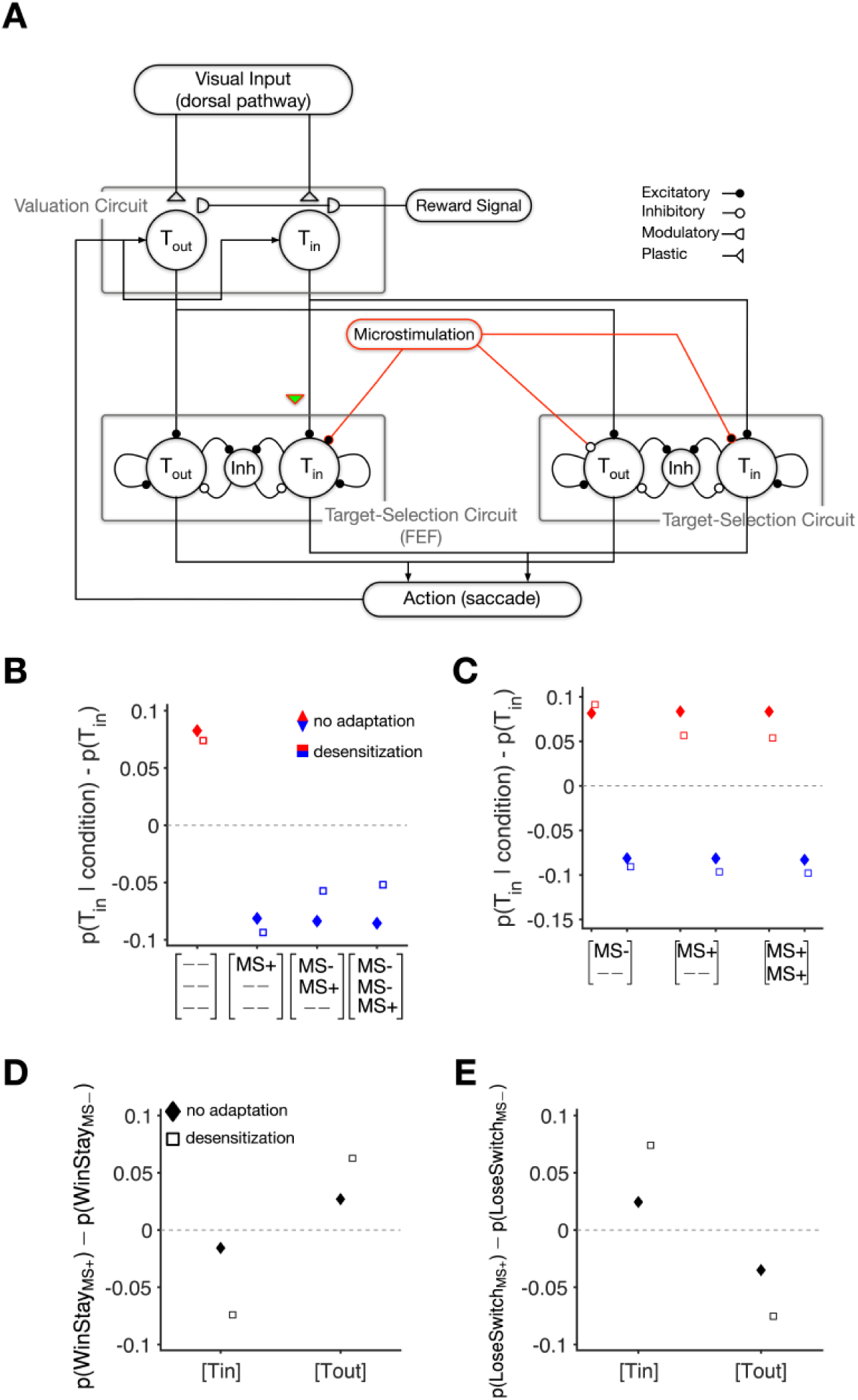
The Schematic and choice behavior of the model with no adaptation and the model with desensitization. (**A**) Both models consisted of three circuits: the valuation circuit and two target-selection circuits corresponding to the FEF and another area in the oculomotor system. The valuation circuit consisted of two pools of value-encoding neurons that were activated upon the presentation of visual targets and projected to the corresponding pools of neurons in the two target-selection circuits. The reward probability associated with two alternative actions was encoded in two sets of plastic synapses onto the value-encoding pools. On microstimulation trials (MS+), the pool of neurons selective for *T_in_* in the FEF target-selection circuit received an extra input. Correspondingly, to simulate cross-area compensation, the value-dependent input to the *T_in_* and *T_out_* pools in the second target-selection circuit were increased and decreased, respectively. In the model with no adaptation, the strength of both the input from microstimulation and the input from the *T_in_* value-encoding pool held constant. In contrast, for the model with desensitization, microstimulation reduced the sensitivity to the input from the *T_in_* value-encoding pool, indicated by a colored triangle. (**B–E**) The choice behavior of the two models in terms of: change in *T_in_* selection on MS+ trials (red) and on MS-trials (blue) that occurred one, two, or three trials following an MS+ trial (B); effects of consecutive microstimulation on target selection (C); and differential response to reward in the presence and absence of microstimulation (D, E). The model with no adaptation is indicated by filled diamonds and model with desensitization is shown with empty squares. Other conventions are the same as in Figure 3. Results are based on 2 × 10^6^ simulated trials to ensure the robustness of the observed effects in the model.

The shortcomings of the above two models suggest that the observed adaptation of the selection bias on MS-trials may require a reward-dependent adaptation of the target-selection network to microstimulation and, thus, an interaction between microstimulation and reward feedback. Therefore, we next asked whether an adaptation of microstimulation effects that depends on the presence or absence of reward could account for our observations. The simulation of this effect was inspired by findings that homeostatic activity regulation, activated by stimulation, can reduce hyperexcitability in brain networks (Chai et al., 2019). However, because mechanisms underlying such homeostatic plasticity are not fully understood, we implemented this effect by creating opposing adaptations in the efficacy of microstimulation and in the strength of input to *T_in_* target (which is endogenous reward) (see **Materials and Methods** for details). We found that a model in which reward produces opposing adaptations in the efficacy of microstimulation and in the strength of endogenous, value-dependent input to *T_in_* target in the FEF target-selection circuit can capture the experimental observations (see **Figure 7A–C**).

Importantly, such opposing adaptations predict that the model should respond to reward and no-reward differently after trials that *T_in_* or *T_out_* was chosen. More specifically, the probability of using the *Win-Stay* strategy on *T_in_* trials (respectively, *T_out_* trials) is predicted to be smaller (respectively, larger) on MS+ trials than on MS-trials (**Fig. 7D**). This model produces such behavior because on MS+ trials the presence of reward increases the efficacy of microstimulation and weakens the efficacy of endogenous reward inputs to the *T_in_* pool in the FEF target-selection circuit. This adaptation results in a decrease in the *Win-Stay* strategy after MS+ trials. However, in the absence of reward, the microstimulation efficacy is reduced following both MS+ and MS-trials (but to a greater extent after MS+ trials) and strengthens the efficacy of endogenous reward inputs to the *T_in_* pool in the FEF target-selection circuit. Therefore, the reward-dependent adaptation does not alter the use of the *Lose-Switch* strategy after MS+ trials (**Fig. 7E**).

Both predictions (or postdictions) were confirmed by our experimental data (**Fig. 3C, D**). First, both monkeys used the *Win-Stay* strategy more often on *T_out_* trials when microstimulation occurred (Wilcoxon ranksum test, *p* = 0.001, *d* = 0.06). In contrast, there was no significant difference in *Win-Stay* on *T_in_* after MS+ and MS-trials (Wilcoxon ranksum test, *p* = 0.22, *d* = 0.02). Second, we found no significant difference in the probability of *Lose-Switch* after MS+ and MS-trials on either *T_in_* or *T_out_* (Wilcoxon ranksum test; *T_in_*: *p* = 0.81, *d* = 0.006, *T_out_*: *p* = 0.95, *d* = 0.002).

Because microstimulation increased *T_in_* selection, the smaller response to reward following a *T_in_* selection with microstimulation indicates an interaction between microstimulation and the reward signals. Similarly, because microstimulation decreased *T_out_* selection, the larger response to reward following a *T_out_* selection with microstimulation points to interactions between the two signals. Importantly, neither the model without adaptation nor the model with sensitization predicted such a pattern of response to reward feedback (**Fig. 6D, E**).

An alternative explanation for the observed interaction between microstimulation and response to reward feedback could be the influence of subjective confidence. Confidence in a choice can affect monkeys’ *Win-Stay* and *Lose-Switch* strategies, and this effect could be different depending on the choice (*T_in_* vs. *T_out_*) and whether microstimulation was applied during a given trial. To test this possibility, we employed two different methods to estimate confidence on each trial (see **Materials and Methods** for details) and examined whether confidence levels could account for the observed response to microstimulation.

**Figure 7.**
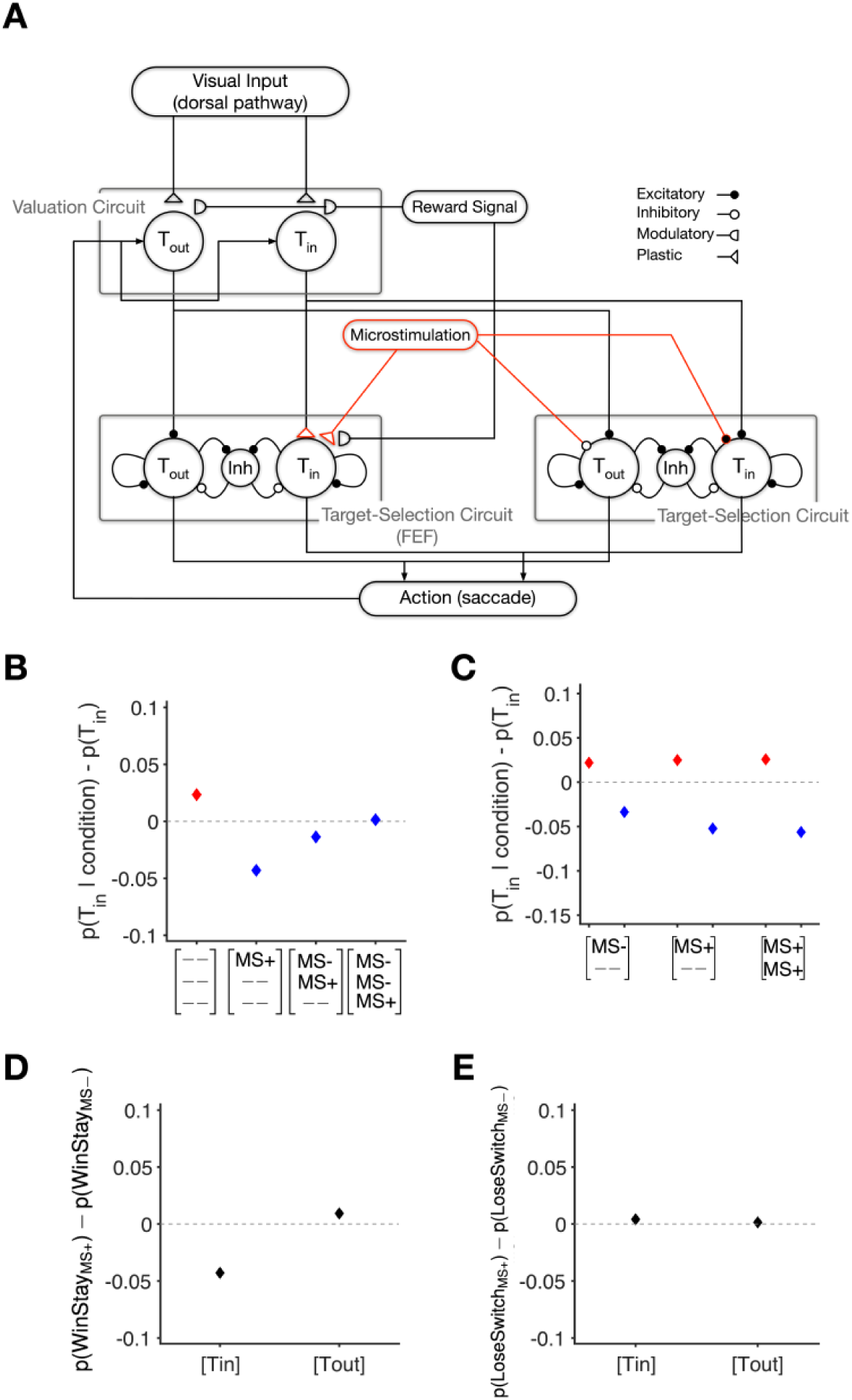
The schematic and choice behavior of the dynamic model with reward-dependent adaptation with opposing adjustments for microstimulation efficacy and endogenous reward inputs to the FEF target-selection circuit. (**A**) The model consisted of three circuits: one valuation circuit and two target-selection circuits corresponding to the FEF and another area in the oculomotor system. The valuation circuit consisted of two pools of value-encoding neurons that were activated upon the presentation of visual targets and projected to the corresponding pools of neurons in the two target-selection circuits. The reward probability associated with two alternative actions was encoded in two sets of plastic synapses onto the value-encoding pools. On microstimulation trials (MS+), the pool of neurons selective for *T_in_*in the FEF target-selection circuit received an extra input. In contrast, the input to the *T_in_* pool in the second target-selection circuit was inhibited. The strength of this input and the input from the *T_in_* value-encoding pool were adjusted by the modulatory reward signal, which resulted in an interaction between the effects of microstimulation and reward (see Methods for more details). (**B–E**) Choice behavior of the model in terms of: change in *T_in_* selection on MS+ trials (red) and on MS-trials (blue) that occurred one, two, or three trials following an MS+ trial (B); effects of consecutive microstimulation on target selection (C); and differential response to reward in the presence and absence of microstimulation (D, E). Other conventions are the same as in Figure 3. Results are based on 2 × 10^6^ simulated trials to ensure the robustness of the observed effects in the model.

After categorizing trials into high-confidence (HC) or low-confidence (LC), we tested whether *Win-Stay* and *Lose-Switch* were different following these types of trials and across the two experimental conditions (i.e., control vs stimulation). We found that in the control condition, the monkeys used *Win-Stay* strategy more often on HC trials, suggesting that the monkeys stayed more on the rewarded choice target when they were more confident about their choices (**Fig. 8A**). Consistently, the monkeys were less likely to switch from the non-rewarded choice target after HC trials (**Fig. 8B**). As expected, these effects were similar for *T_in_* and *T_outt_*, as there was no stimulation in the control condition.

**Figure 8.**
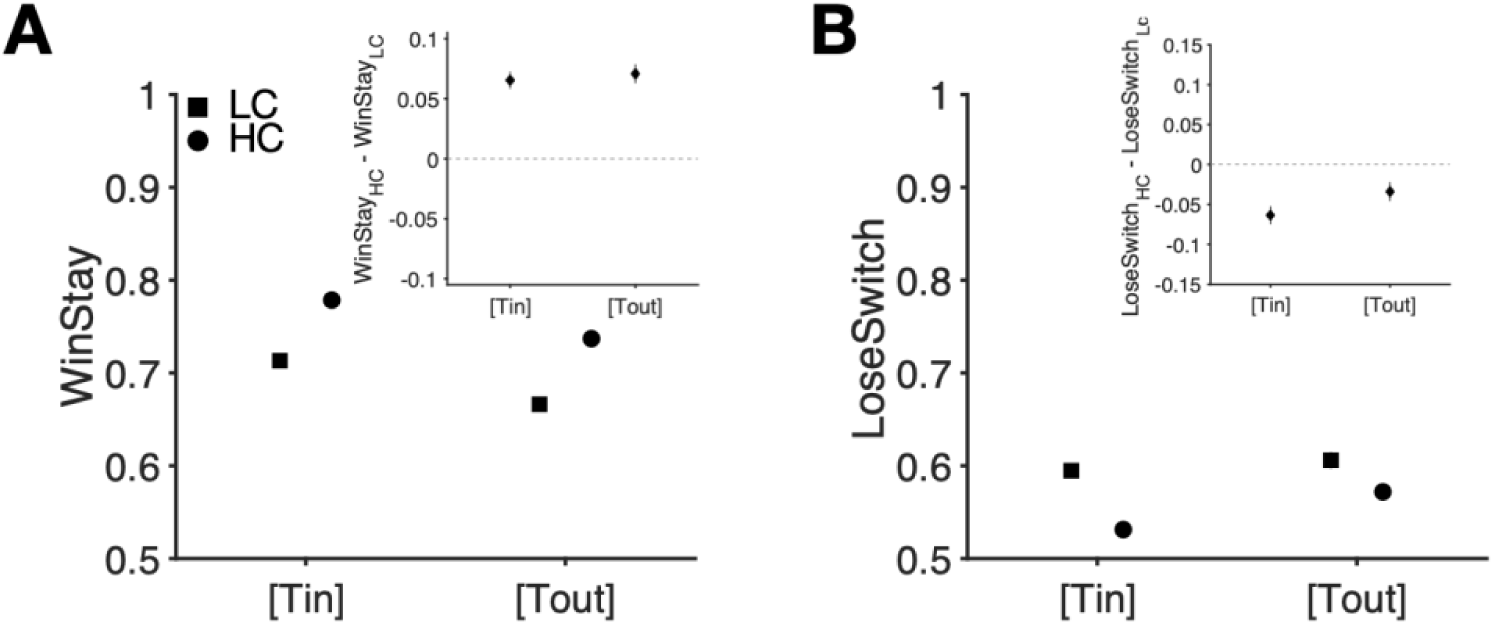
Dependence of *Win-Stay* and *Lose-Switch* strategies on confidence during control sessions. High-confidence (HC) and low-confidence (LC) trials are defined using median reaction time. (**A**) Plot shows the probability of *Win-Stay* separately for HC (circle) and LC (square) trials and for the *T_in_* and *T_out_* targets. The inset depicts the difference in Win-Stay strategy between HC and LC trials. (**B**) Plot shows the probability of *Lose-Switch* separately for HC and LC trials and for the *T_in_* and *T_out_* targets. The inset depicts the difference in *Lose-Switch* strategy between HC and LC trials.

As mentioned above (**Fig. 3C, D**), the *Win-Stay* and *Lose-Switch* during the stimulation condition were influenced by both stimulation and the choice such that the monkeys stayed less on *T_in_* if it was selected and rewarded and there was a stimulation compared to when there was no stimulation. In contrast, the monkeys stayed less on *T_out_* if it was selected and rewarded and there was *T_in_* stimulation compared to when there was no stimulation. Interestingly, we found that the pattern of *Win-Stay* was similar across both HC and LC trials, emphasizing the consistency in behavior regardless of confidence level (**Fig. 9A–B**). Moreover, the frequency of *Lose-Switch* strategy did not depend on the combination of stimulation and choice, regardless of the confidence level (**Fig. 9C–D**). Overall, results of confidence analyses suggest that although confidence affects monkeys’ response to reward feedback, variations in confidence levels on rewarded and unrewarded trials cannot account for our experimental findings, which appear to be better accounted for by the interaction between reward and microstimulation.

**Figure 9.**
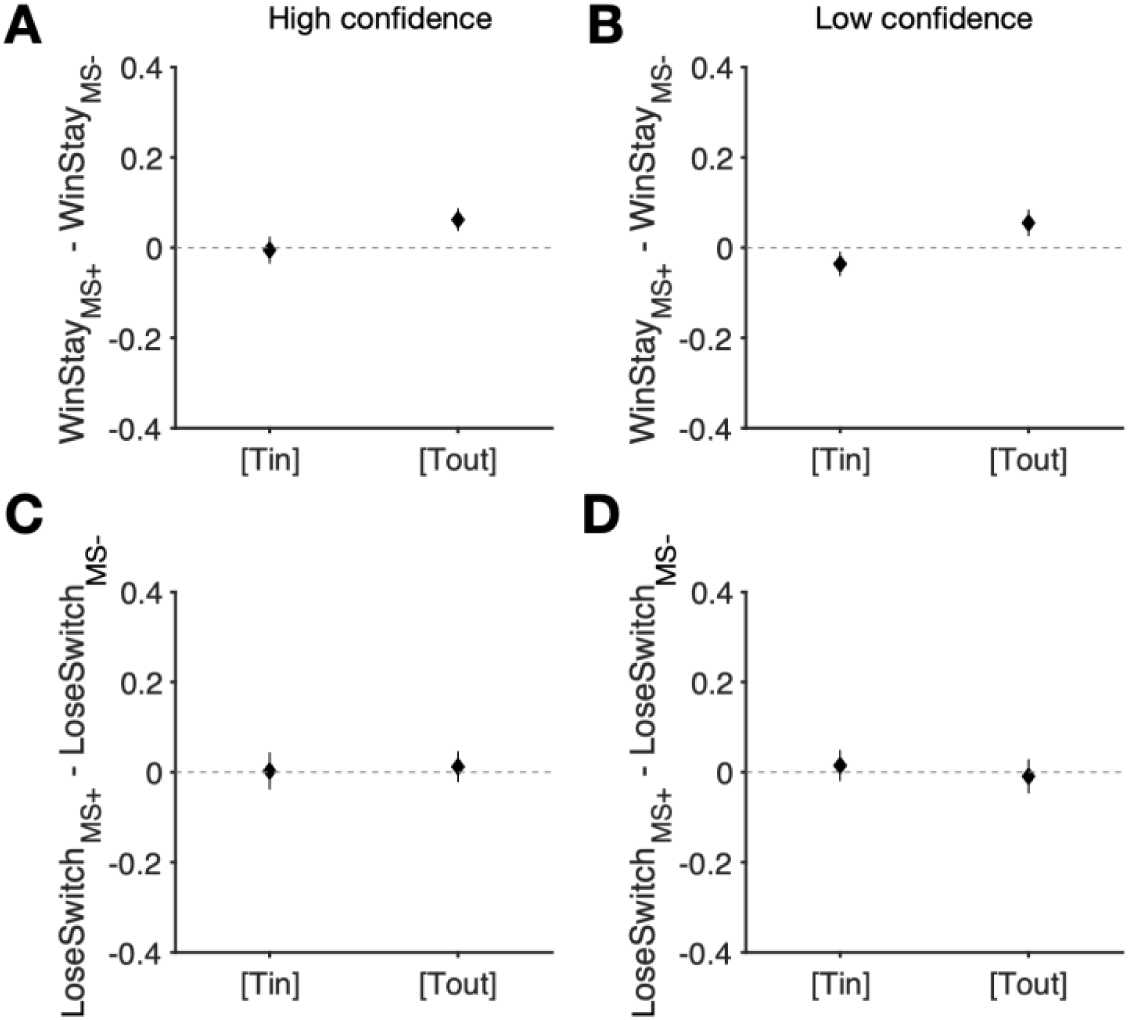
Difference in *Win-Stay* and *Lose-Switch* strategies between stimulated and non-stimulated trials was not affected by confidence level. (**A**) Plot shows the difference in *Win-Stay* is plotted in HC trials in which microstimulation occurred (MS+) and the trials in which no microstimulation (MS-) occurred. (**B**) Plot shows the difference in *Win-Stay* is plotted in LC trials in which microstimulation occurred (MS+) and the trials in which no microstimulation (MS-) occurred. (**C–D**) Similar to panels A and B, but for *Lose-Switch*.

Finally, we asked whether the cross-area compensation was essential for replicating the experimental results. To that end, we created three models with a single target-selection circuit, even though such models are not consistent with existing knowledge of the oculomotor system. More specifically, we simulated three models: a base model with no adaptation to microstimulation, a model with desensitization, and a model with reward-dependent adaptation (see **Materials and Methods** for more details). Interestingly, we found that only the model with reward-dependent adaptation was able to replicate the pattern of our experimental data (**Fig. 10**). These results suggest that the interaction between reward and microstimulation through reward-dependent adaptation is a critical mechanism for capturing the experimental data. However, the presence of a second target-selection circuit remains a necessary component for a biologically plausible network model. Together, our experimental and modeling results provide strong evidence that robust saccadic choice relies on the interaction between adaptation to microstimulation and endogenous reward signals within the oculomotor systems and related circuits.

**Figure 10.**
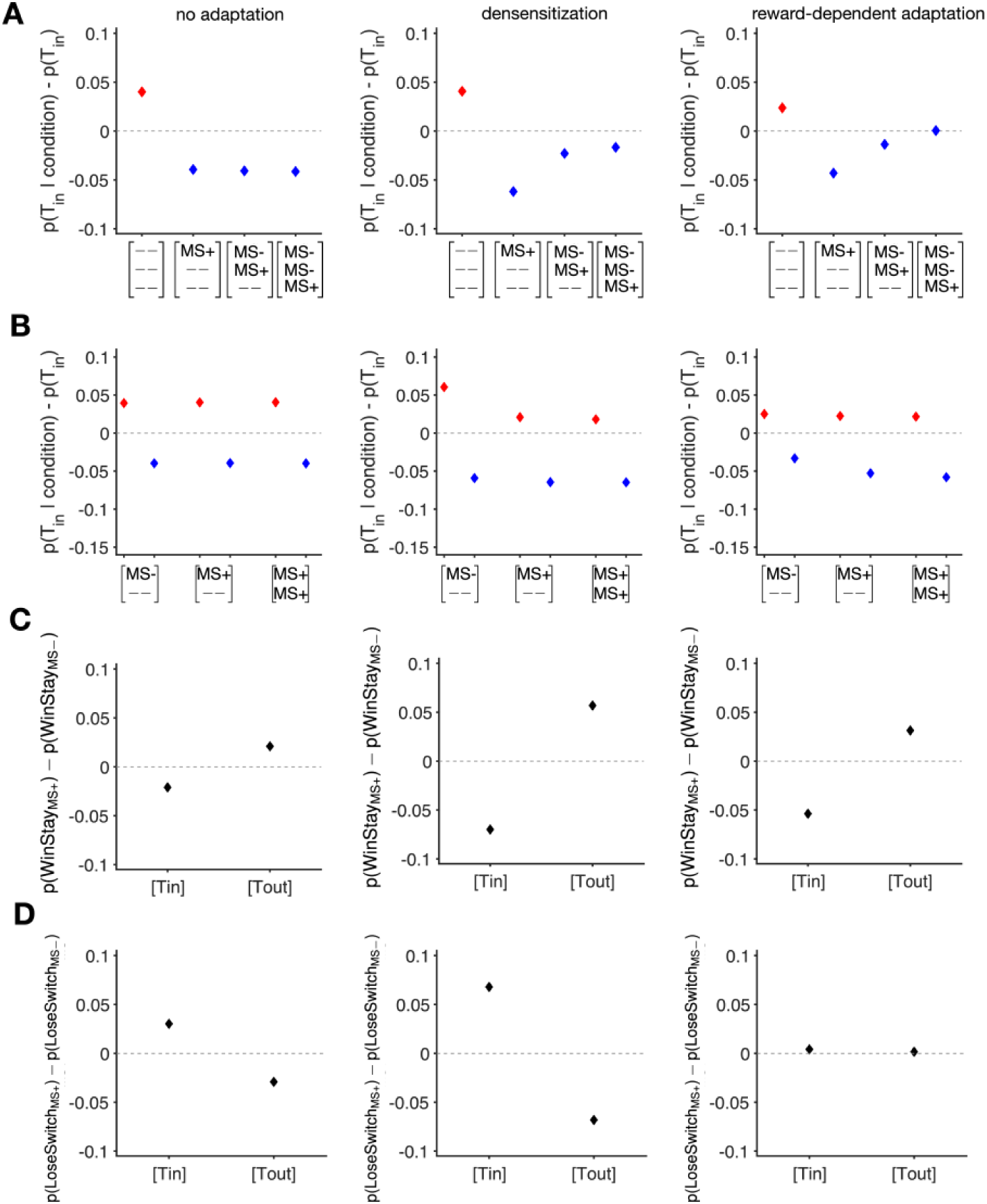
Behavior of models with a single target-selection circuit. Plots across rows A–D capture: change in *T_in_* selection on MS+ trials (red) and on MS-trials (blue) that occurred one, two, or three trials following an MS+ trial (A); effects of consecutive microstimulation on target selection (B); and differential response to reward in the presence and absence of microstimulation (C–D). Plots across columns show the choice behavior of the model with no adaptation to microstimulation (left panels), the model with desensitization to microstimulation (central panels), and the model with reward-dependent adaptation to microstimulation (right panels). Conventions are the same as in Figures 6–7. The choice behavior of these models is qualitatively similar to their corresponding models that consider cross-area compensation. These results suggest that the interaction between reward feedback and microstimulation is essential in replicating the monkeys’ choice behavior. Results are based on 2 × 10^6^ simulated trials to ensure the robustness of the observed effects in the model.

## Discussion

By applying microstimulation during a dynamic value-based learning task, we found a specific pattern of change in choice behavior in response to microstimulation and reward feedback. More specifically, subthreshold electrical microstimulation of the FEF increased the selection of a target within the RF (*T_in_*) on a trial-by-trial basis. On non-stimulated trials following microstimulation, we found a bias away from *T_in_* and this bias decreased as a function of the time since the last stimulation. The bias toward *T_in_* did not increase with consecutive stimulation, but the opposite bias on the subsequent non-stimulated trials did.

It is possible that the observed pattern of response to microstimulation was partially influenced by the dynamic reward schedule used in our experiment, where the probability of reward on subsequent selection of the same target gradually decreased. However, similar long-lasting effects of microstimulation on non-stimulated trials have been previously reported for microstimulation of SC during a task in which the statistics of correct response were manipulated across a block of trials (Crapse et al., 2018). In addition, simulation results from the model with no adaptation demonstrate that reward feedback alone cannot fully explain our experimental findings. This suggests that the absence of an overall bias toward *T_in_* is not merely a result of reward feedback promoting the monkeys to compensate for biases caused by microstimulation. Critically, we found that microstimulation and reward signals interacted to guide saccadic choice. More specifically, the probability of choosing the target rewarded in the preceding trial depended on the presence or absence of the stimulation in that trial. This observation and the effect of consecutive microstimulation on the subsequent non-stimulated trials point to an adaptive, reward-dependent compensatory mechanism within the oculomotor system and related circuits that allow for robust saccadic choice behavior.

The network model that best captured our experimental results included a competitive mechanism for adjustment of microstimulation efficacy and reward input in the oculomotor systems and related circuits. This suggests that microstimulation signals into the FEF compete with signals from the endogenous reward system, allowing each pathway to be upregulated or downregulated at the expense of the other. Moreover, this model can account for general characteristics of the effects of external perturbations on behavior and explains some of the more idiosyncratic effects of microstimulation in other studies. For example, Ni and Maunsell (Ni & Maunsell, 2010) found that monkeys could be trained to detect microstimulation of V1 corresponding to particular retinotopic locations. Surprisingly, however, training to detect V1 microstimulation caused a profound impairment in the monkeys’ ability to detect veridical visual stimuli at the same visual location, and re-training on visual detection impaired the detection of microstimulation. Although we observed adaptation on the order of several trials rather than tens or hundreds of trials, as in Ni and Maunsell (Ni & Maunsell, 2010), our model is able to explain their results qualitatively.

The proposed adaptive and compensatory mechanism due to the interaction between microstimulation and reward signals within the oculomotor system can provide an alternative explanation for inconsistent effects of electrical microstimulation on choice behavior or, equivalently, how the oculomotor system is robust against internal and external perturbations. For example, a recent meta-analysis study on the utilization of electrical stimulation for cognitive therapy (Grover et al., 2023) concluded improvements in working memory and attention with electrical stimulation. In contrast, they found that electrical stimulation did not cause significant modulations of cognitive functions such as motor learning and decision making. Our results are consistent with these findings because goal-driven and reward-dependent processes involved in learning and decision-making tasks are likely to be more robust to electrical stimulation due to stronger interaction between stimulation and reward signals during those tasks.

Our implementation of a homeostasis-like mechanism activated by microstimulation sheds light on the paradoxical observation that electrical stimulation can reduce hyperexcitability in brain networks, a phenomenon associated with various neurological disorders such as epilepsy and neuropathic pain (Chai et al., 2019). Interestingly, both electrical stimulation and medications that inhibit neuronal activity have been found to reduce this hyperexcitability effectively. This suggests that the increased excitatory currents resulting from electrical stimulation may be counteracted by further adaptations within the same circuits. Indeed, several studies have proposed that the therapeutic effects of stimulation in hyperexcitability-related neural disorders are attributed to homeostatic activity regulation resulting from the stimulation (Chai et al., 2019).

Finally, our findings of reward-dependent, adaptive and compensatory mechanisms in target selection underscore the need for caution in understanding and interpreting the results of studies that employ stimulation for neurorehabilitation following brain damage (Cappon et al., 2016; Dolbow et al., 2014; Plow et al., 2009), use stimulation to modulate cognitive processes such as memory (Aaronson et al., 2021; Alekseichuk et al., 2016; Antonenko et al., 2013), attention (Clayton et al., 2018; Dallmer-Zerbe et al., 2020), executive control (Borghini et al., 2018; Bramson et al., 2020), motor learning (Giustiniani et al., 2019; Harada et al., 2020), and learning and decision making (Sela et al., 2012; Zavecz et al., 2020), or to treat depression (Shekelle et al., 2018). More specifically, the majority of stimulation protocols involve participants receiving reward feedback, which can be explicit, like reward points, or implicit, such as a correct/incorrect message displayed on the screen. As demonstrated in our findings, such feedback can competitively interact with stimulations, systematically reducing their overall effect on behavior. Moreover, the individual variability observed in our study suggests that the interaction between stimulation and reward feedback may lead to more idiosyncratic outcomes, making the interpretation of these results specific to each individual. These indicate that to accurately identify the effects of stimulation, both local and global impacts of stimulation must be analyzed and understood in relation to reward outcomes on specific stimulation trials.

## Acknowledgments

We thank Chanc VanWinkle Orzell for her helpful comments on the manuscript. This work is supported by the National Science Foundation (CAREER Award BCS1943767 to A.S.) and the National Institutes of Health (NIH EY014924 to T.M.).

